# DPSleep: Open-Source Longitudinal Sleep Analysis From Accelerometer Data

**DOI:** 10.1101/2021.02.02.429455

**Authors:** Habiballah Rahimi-Eichi, Garth Coombs, Constanza M. Vidal Bustamante, Jukka-Pekka Onnela, Justin T. Baker, Randy L. Buckner

## Abstract

Wearable devices are now widely available to collect continuous objective behavioral data from individuals and to measure sleep. Here we introduce a pipeline to infer sleep onset, duration, and quality from raw accelerometer data and then quantify relationships between derived sleep metrics and other variables of interest. The pipeline released here for the deep phenotyping of sleep, as the “DPSleep” software package, uses (a) a stepwise algorithm to detect missing data; (b) within-individual, minute-based, spectral power percentiles of activity; and (c) iterative, forward- and backward-sliding windows to estimate the major Sleep Episode onset and offset. Software modules allow for manual quality control adjustment of derived sleep features and correction for time zone changes. In this report, we illustrate the pipeline with data from participants studied for more than 200 days each. Actigraphy-based measures of sleep duration are associated with self-report rating of sleep quality. Simultaneous measures of smartphone use and GPS location data support the validity of the sleep timing inferences and reveal how phone measures of sleep timing can differ from actigraphy data. We discuss the uses of DPSleep in relation to other available sleep estimation approaches and provide example use cases that include multi-dimensional, deep longitudinal phenotyping, extended measurement of dynamics associated with mental illness, and the possibility of combining wearable actigraphy and personal electronic device data (e.g., smartphone, tablet) to measure individual differences across a wide range of behavioral variation in health and disease.

---

While the human behavioral repertoire varies in many respects between individuals, prolonged daily episodes of sleep behavior are expressed nearly ubiquitously in all members of our species, as they are innate and undergird both physical and mental health across the lifespan. Multiple studies suggest that sleep loss or poor sleep quality are predictors (and potentially moderators and/or mediators) of mental illness symptoms and poor cognitive performance (Maglione et al., 2014; Phillips et al., 2017; Blake, Trinder, and Allen, 2018; Snyder et al., 2018; André et al., 2019; Merikangas et al., 2019). Sleep timing in modern society is affected by interactions with digital devices that enable low-burden approaches to measure sleep (Adams, Daly, and Williford, 2013; Demirci, Akgonul, and Akpinar, 2015; Cabré-Riera et al., 2019; Tashjian, Mullins, and Galván, 2019). Sleep timing patterns can be measured over extended periods of time to provide a window into the rhythms and dynamics of a person’s life. In this paper we describe an open-source sleep analysis pipeline called DPSleep, referring to the deep phenotyping of sleep, that we offer to the community as a platform to facilitate longitudinal studies of sleep using data from widely available wearable devices (https://github.com/harvard-nrg/dpsleep-extract).

The current gold standard for documenting sleep timing and content is polysomnography (PSG), during which multiple physiological measures are recorded, usually in a clinic or laboratory setting (Sadeh, 2011a; Hirshkowitz, 2017) or recently in ambulatory settings (Fonareva et al., 2011; Smith et al., 2020). Although PSG, including its multiple physiological measures, is a comprehensive assessment of sleep stages, there are limitations associated with cost and subject burden, and it is difficult to obtain in-patient or at-home versions of PSG for extended time periods (Newell et al., 2012; Kaplan et al., 2017). Actigraphy has been suggested as an efficient and reliable alternative to measure certain features of sleep patterns in natural, at-home settings (Ancoli-Israel et al., 2003; Sadeh, 2011b; Stone and Ancoli-Israel, 2017). Actigraphy data, estimated from accelerometers, is a common output of many wearable and held devices including wrist watches, ankle and wrist bands, smartglasses, sewn-in or attached devices, and smartphones (Schenk et al., 2011; Evenson, Goto, and Furberg, 2015; Staples et al., 2017; Wright et al., 2017; Scott, Lack, and Lovato, 2020; Smith et al., 2020). Thus, actigraphy measurement provides a means to study sleep patterns at an unprecedented scale. At the same time, openly available feature-extraction algorithms with capability to retain and present the features from raw to derived measures are essential for reproducible large-scale understanding of human sleep, even in the presence of the proprietary algorithms associated with many of the devices.

In actigraphy-based sleep assessment using watch-like wristbands, acceleration is typically measured in three dimensions; some devices also measure ambient light, temperature, heart rate, or electrodermal activity (EDA) (Bianchi, 2018). A body of literature investigating the sensitivity and specificity of wristbands to detect sleep parameters including total sleep time (TST), sleep onset latency (SOL), wake after sleep onset (WASO), and sleep efficiency (SE) has evolved (Soric et al., 2012; Shin, Swan, and Chow, 2015; Madrid-Navarro et al., 2019). While several of these approaches have been validated using PSG and self-report sleep quality ratings under certain controlled conditions, there is ongoing research to validate the software and develop in-house algorithms for different applications, and under real-world situations (Cellini et al., 2016; Mantua, Gravel, and Spencer, 2016; John et al., 2019). Several of the devices, in order to save memory and battery, provide pre-processed one-minute averaged acceleration data (Cole et al., 1992; Bakken et al., 2014; Meltzer et al., 2015; Bellone et al., 2016; Merikangas et al., 2019) while others provide continuous high-frequency data. (Dillon et al., 2016; Crowley et al., 2019; Jones et al., 2019).

The present study contributes to this evolving field by providing a comprehensive pipeline to analyze raw accelerometer data to estimate minute-based activity and detect the major Sleep Episode, defined as the longest continuous sleep episode of at least 100 minutes. The overall goals were to (1) develop an open-source processing pipeline for commonly available accelerometer data to detect major Sleep Episodes, (2) apply and validate the estimation procedure using real-world data including individuals studied over extended time periods who independently rated their sleep, and (3) apply the processing pipeline to exemplar data to illustrate its application. For this final goal, we analyzed data from undergraduate students who were studied over 6-9 months during college to illustrate sleep patterns that fluctuate with environmental demands and in relation to other self-report measures of sleep and mental state. We also analyzed data from two individuals who were outpatients living with severe mental illness to illustrate the boundaries of the methods and their ability to measure dramatically altered sleep patterns.

## Materials and Methods

### Participants

Participants were enrolled in two distinct cohorts to obtain actigraphy data across a range of subjects. We describe the samples separately.

#### Study 1: Undergraduate Study

Six undergraduate participants (all 19 years old; 3 females) were recruited from a local private institution and participated for one academic year, including a buffer extending into summer break. These individuals had successfully participated in a shorter, earlier pilot study that did not use the present actigraphy device or processing pipeline and were enrolled here to generate extended data for the present purposes collected over 165-268 days. Participants were compensated per hour for the lab visits and for completing online daily questionnaires and given milestone bonuses to encourage continued participation. Participants were required to be enrolled full-time in classes and own an iPhone or Android smartphone compatible with the study smartphone application, Beiwe, which is part of the open-source Beiwe platform for digital phenotyping (Torous et al., 2016). The Beiwe application was configured to collect passive phone use, phone acceleration, and GPS data at an almost continuous rate as well as active self-report data on a regular, daily basis. Since each participant serves as her or his own baseline, participants were not excluded for current or past psychiatric disorders or medication use. All study procedures were approved by the Institutional Review Board of Harvard University.

#### Study 2: Clinical Study

Two individuals (ages = 62, 24; 1 female) were recruited from an ongoing cohort following the clinical progression of severe mental illness at a local hospital. Individuals were diagnosed with psychotic disorders (bipolar, n=1; schizophrenia, n=1) using the structured clinical interview for DSM-IV (SCID, First et al., 2007). Participant enrollment for this study targeted obtaining >1 year of data for each participant (durations: 543, 309 days), which included nearly continuous collection of smartphone and actigraphy data via the Beiwe platform (Torous et al., 2016) and wearable watch, respectively, from each participant. Participants were compensated for these data, as well as for monthly in-person study visits during which clinical assessments were recorded to quantify disease progression using clinical gold standard measures. Milestone bonuses were also provided to encourage continued participation. All study procedures were approved by the Institutional Review Board of Partners Healthcare.

### Wrist Actigraphy and Ancillary Data Acquisition

The present pipeline was developed using tri-axial acceleration data from a commercially available waterproof watch worn on the wrist (GENEActiv; Activinsights; Kimbolton, UK) and is intended as a general open resource for processing accelerometer data from any device saving raw tri-axial, high-frequency, continuous accelerometer data, sampled at a fixed and known rate. Missingness of data is assumed to occur completely at random. Data saved as minute-based or shorter activity estimates can also be accommodated. The frequency of data sampling was set to 30Hz for Study 1 and 20Hz for Study 2. Following the initial consent and receipt of the watch, individuals in Studies 1 and 2 visited the lab every 4-5 weeks to return the watch and receive a new fully charged watch with formatted memory; participants in Study 2 were given the watches during their in-patient study. Participants were instructed to wear the watch continuously including during sleep and when bathing. Using the same sampling rates across modalities of 30Hz in Study 1 and 20Hz in Study 2, the watch collected acceleration (g), light (lux), and ambient temperature (°C). In addition, the wristband recorded key presses. Subjects were instructed to press the key when they started to go to sleep and also when they woke in the morning. The acceleration data are the primary data used for automatic detection of episodes that would be scored as sleep with additional corrections from light and key press data when available and when necessary.

Data obtained from a smartphone (iPhone or Android) were used as an ancillary data source (Onnela and Rauch, 2016; Torous et al., 2016). None of the automated processing and manual Sleep Episode adjustments used data from the smartphone. The smartphone data provided valuable independent information for validation. Individuals installed the research smartphone application, Beiwe, to collect active (questionnaire), passive phone use (via timestamping of lock-unlock events), accelerometer, and Global Positioning System (GPS) data (Torous et al., 2016; 2017; Barnett et al., 2018). GPS location of the phone was sampled every 10 min for a 2-min duration, and phone acceleration was sampled 10 sec on, 10 sec off. At 5PM each day, an in-app questionnaire appeared that asked about the quality of the previous night’s sleep on a Likert scale (from Terrible to Exceptional), the amount of caffeine consumed in the prior 24 hours (from None to Five+ drinks) (see Appendix I), and a set of questions about their mood and social and academic activities. Answers to these questions and passive estimates of phone use were used to validate the identification of the major Sleep Episode from the independent watch actigraphy data.

### DPSleep: A Processing Pipeline for Deep Phenotyping of Sleep

#### Raw actigraphy data and removal of missing (wrist-off) data

Raw wrist actigraphy data are saved originally as large, compressed files each containing multiple weeks of data. Each file comprises a table with columns of acceleration, light, and temperature (depending on the device). To overcome the challenge of time-consuming access to specific rows during analysis, the first step in our processing pipeline is to parse and save the data into separate daily files from 12:00AM to 11:59PM. For days on which a watch change occurred (to allow continuous data sampling), the new, already charged watch was placed on the wrist. Since the old watch keeps collecting non-wrist data until it is connected back to the data extraction station, the data from the new watch were designated to overwrite any data from the previous watch at the same clock times.

Raw actigraphy data (30Hz) are displayed for one daily data file illustrating each separate accelerometer trace (Figure 1). This type of raw actigraphy data measures acceleration and are thus most sensitive to dynamic movements (Yang and Hsu, 2010). The amplitude of the fluctuations, measured in units of gravity, g, reflect the relative acceleration of the actigraphy device in relation to the gravity of Earth. Each axis – x(g), y(g), z(g) – reflects linear acceleration along one of the device’s three orthogonally-positioned accelerometers. The raw data always includes the Earth gravity component, which can be reflected on different axes depending on the orientation of the device. The separate channels are highly correlated and, for our purposes here, provide redundant information that can be interrogated separately or combined to make estimates more robust. The amplitude of the accelerometer fluctuations is variable across the awake hours but shows a stark reduction of fluctuations, in all three axes, during the sleep period. As will be targeted below, the DPSleep processing pipeline is optimized to detect the reduction in fluctuations and estimate a single extended episode each day.

**Figure 1.**
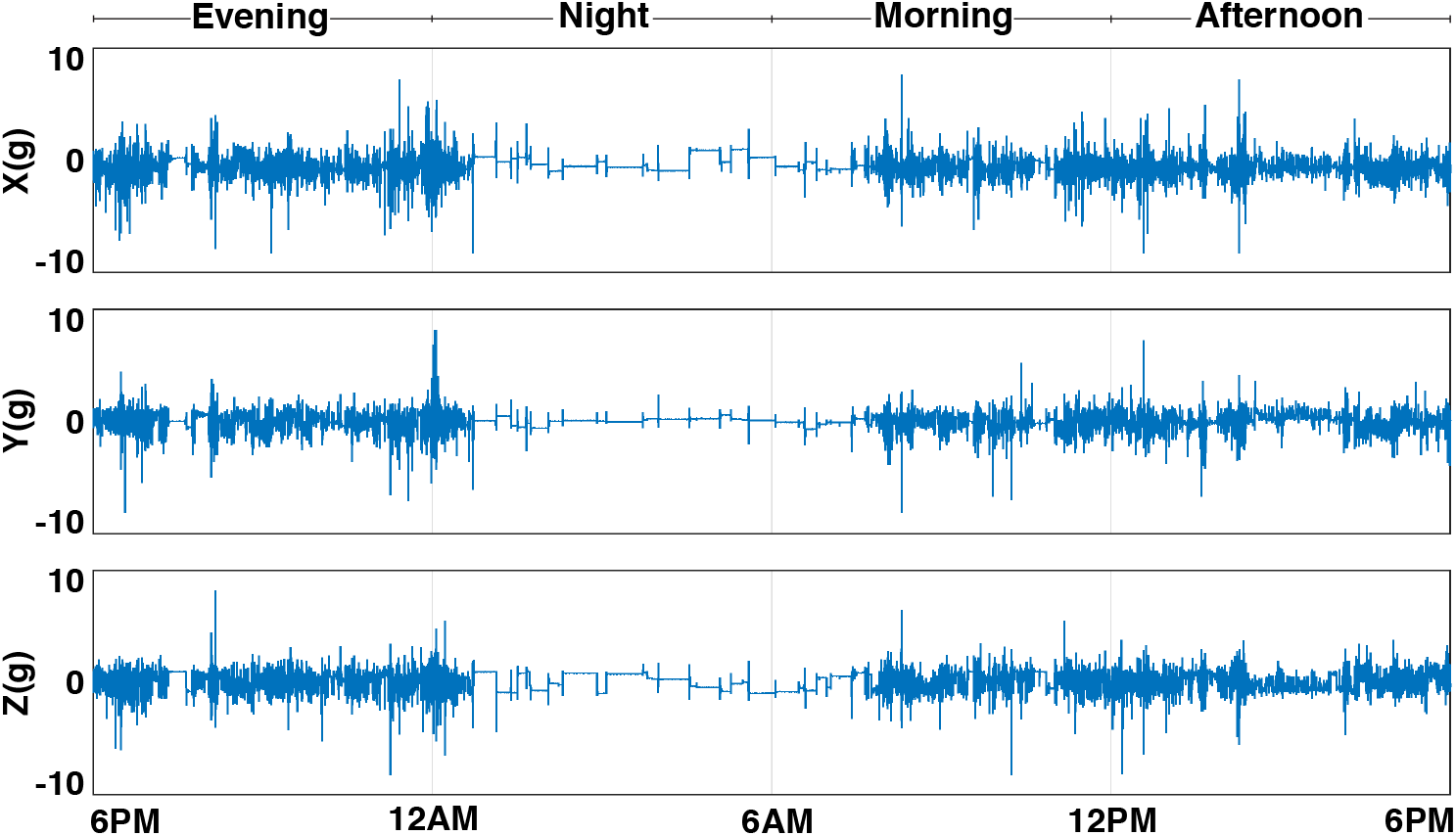
Three-axis accelerometer data for a day of data. The raw acceleration signal along three axes sampled here at 30Hz and scaled to 9.8 m/s^2^ gravity intuitively shows the change in the dynamics of the signal, including the Sleep Episode from approximately 12AM to 7AM.

An immediate challenge is that, even for compliant subjects with charged, waterproof actigraphy watches, the subjects occasionally remove their devices. Figure 2 shows an epoch where the watch is off the wrist. Unlike the Sleep Episode that contains residual periodic low-levels of fluctuations, the accelerometer traces are nearly flat during the wrist-off periods. The first step is to detect and remove the wrist-off minutes. Since the focus of these analyses is on identifying the major Sleep Episode, a window size of 150 minutes is used. The standard deviation of the acceleration time series for each of the three axes are calculated for each minute. A forward and backward moving window of standard deviation for each axis is calculated for window size of 150 minutes. Then, the Root Mean Square (RMS) of the moving average standard deviation values for three axes is compared to a small experimental threshold of 0.0185 to detect the wrist-off minutes. This threshold was determined based on more than 600 hours (five lab volunteers for 5-7 days) of annotated data collected in-house. The minute of data is considered wrist-off if the RMS of the three channels’ variance shows a value below this threshold. The remaining data are considered for further analysis.

**Figure 2.**
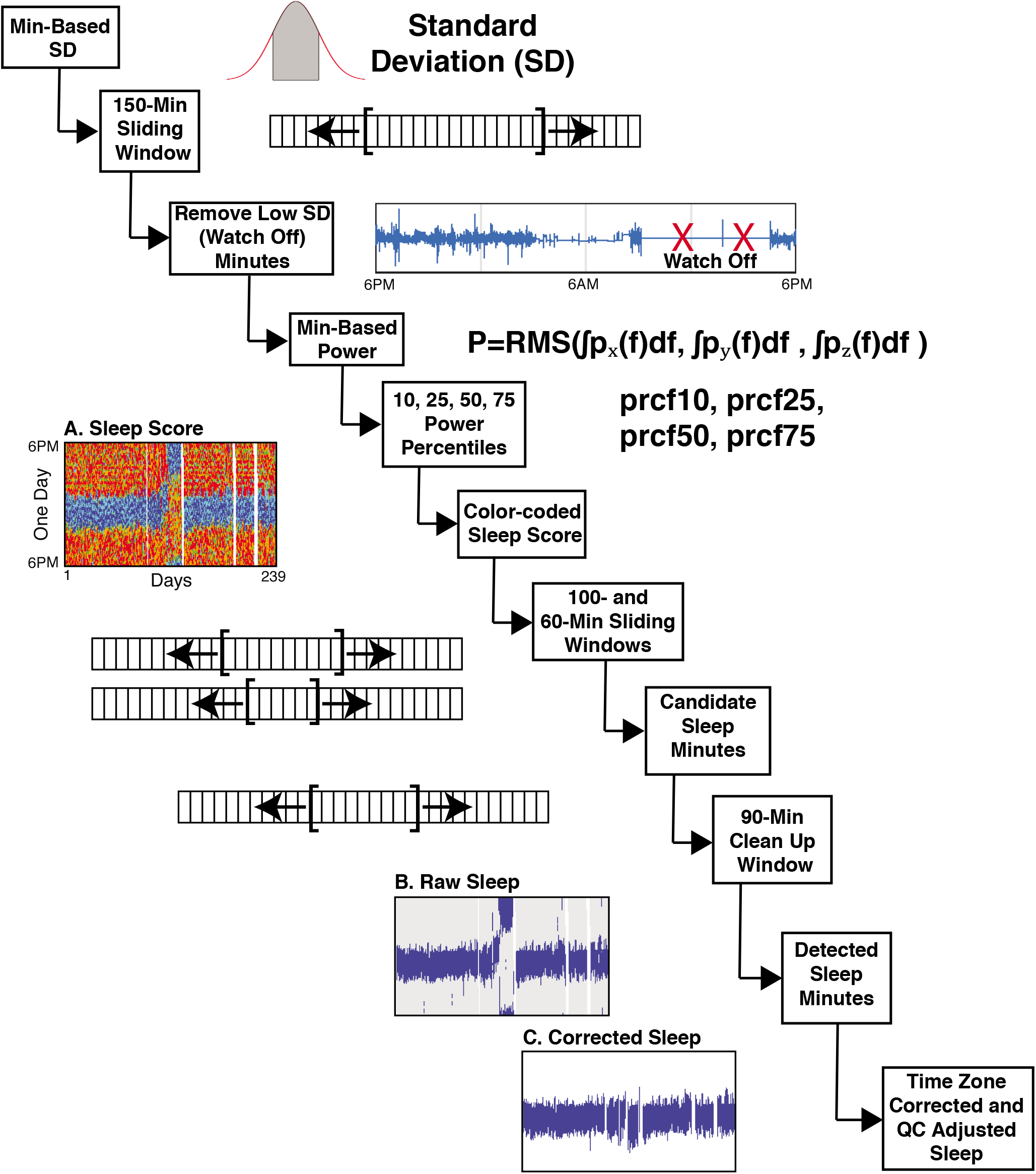
The DPSleep processing pipeline. The sequential steps in the processing pipeline are illustrated with example data for key steps. The pipeline begins with raw accelerometer data and derives estimates of individualized sleep scores (A) as well as raw (B) and corrected Sleep Episodes (C). Output variables in Table 1 are calculated based on the corrected Sleep Episodes as estimated in C. Specifically, as the first step, the standard deviation of the acceleration along three axes is averaged over 150-minute windows forward and backward to find the wrist-off minutes. The Welch equation is then used to calculate the power density spectrum of the acceleration signal at different frequencies, and the area under the curve estimates power as the root mean squared integrated over the three axes. Minutes are classified based on 10, 25, 50, and 75 percentile thresholds and can be visualized (A; blue, cyan, green, orange, and red display the minutes based on increasing power scores). A series of forward and backward moving average windows are used to flag the candidate Sleep Episodes that are then further filtered to derive the raw sleep estimate (B). Quality control and adjustment for the local time zone then yield to the final corrected sleep estimate (C).

#### Scoring activity level for each minute

After removing the wrist-off minutes from the analysis, the power density spectrum of the acceleration signal in each minute is calculated using Welch’s formula (Welch, 1967). The power of the signal in each minute is the area under the curve of the power density spectrum. Within-individual power thresholds are then determined for 10, 25, 50, and 75 percentiles. A 15-min wide window of the average RMS power that combines across the three axes is used to find the percentile thresholds specific to each individual. Data from each minute of the study are classified based on their spectral power in comparison with these percentile cut-offs and color-coded daily maps of the activity scores for each individual are generated (Figure 2, center left). Highly active minutes are colored in red and orange, minutes with medium activity in green, and low active minutes in blue and cyan. The daily maps provide intuitive information about the sleep pattern of the individual. Since the watch is recording continuous actigraphy without reference to the external world, when the time zone is changed due to traveling, the sleep pattern is shifted and will need to be accommodated at a later stage of processing, as can be seen in this figure at approximately day 150. Standard/ Daylight Saving Time transitions similarly require correction.

**Table 1.**
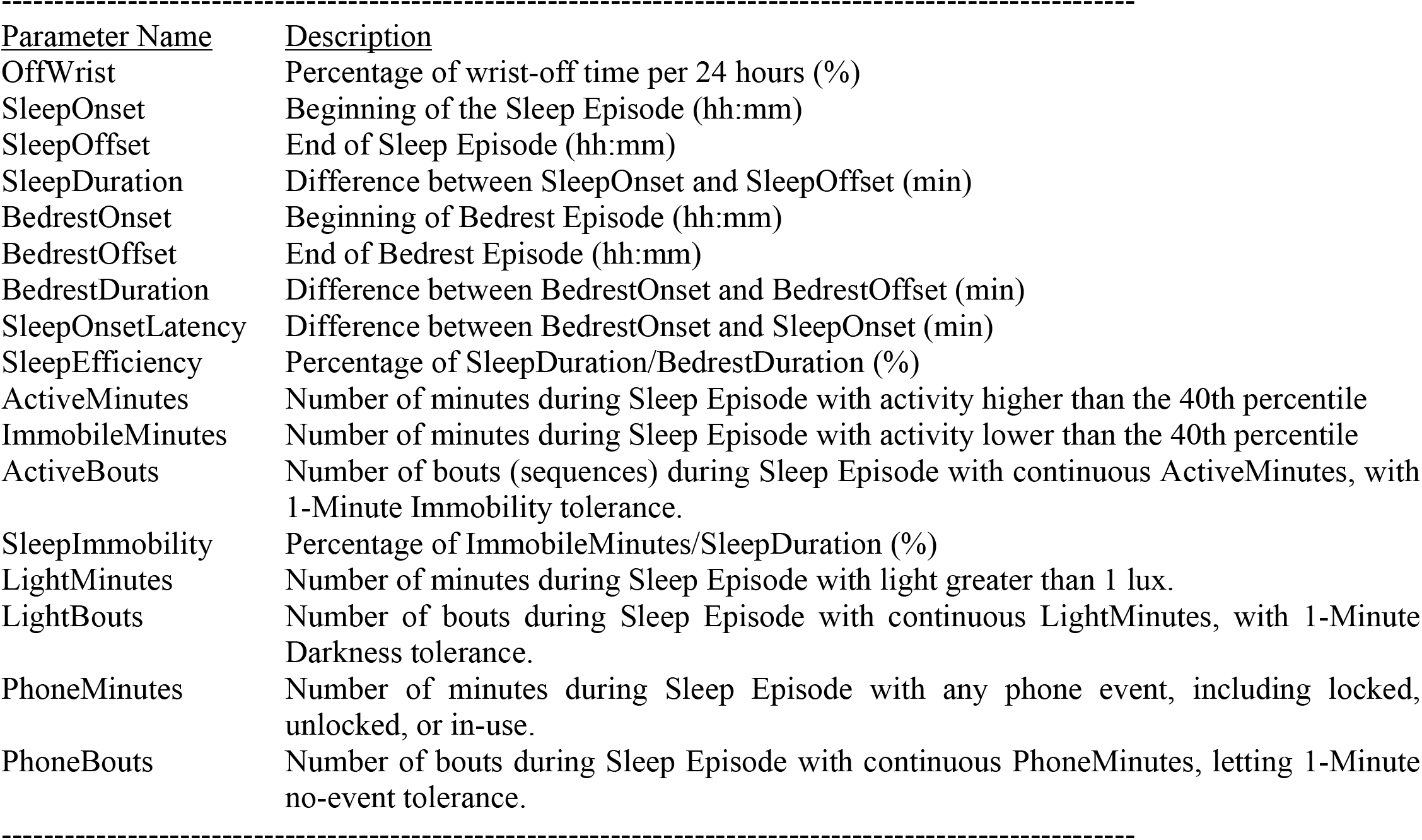
Sleep Variables Generated by the DPSleep Pipeline

#### Estimating the major Sleep Episode

To automatically estimate the timing of the Sleep Episode, we utilize multiple moving windows that slide over a weighted transformation of the minute-based activity levels. The minutes with less than 25% activity (25^th^ percentile of the empirical distribution of activity for the person) are assigned a sleep score of 1 and those with higher than 50% activity are penalized by negative experimental scores (−0.75 and −1). Then, two moving windows of narrow (60-min) and wide (100-min) sizes are used to sweep the scores and find a provisional nocturnal episode of sleep while the narrower window adjusts the beginning and the end estimates of the major Sleep Episode. The separation of task between two window sizes was found, in pilot analyses, to better capture the distinct targeted events, where detecting the nocturnal Sleep Episode benefitted from the larger window, but the precision of the sleep onset and offset time estimates benefitted from the smaller window. A ‘clean-up’ 90-min wide moving window was then used to connect adjacent short candidate Sleep Episodes with more than three-quarter sleep-scored minutes in each 90-min window. However, this process, on some occasions, still left two separate candidate Sleep Episodes that were separated by a period of activity during the middle of the night. As a convention, the automated algorithm joined discontinuous Sleep Episodes into one longer episode if they fell within 22.5 minutes (a quarter of the 90-min window) of one another. This is a decision of convention and occurred in 2.3% of the cases in which sleep was measured in healthy young adults.

The outcome of these steps is an estimate of a single provisional Sleep Episode for each day. The estimate, via the filtering approaches used, usually underestimates the full Sleep Episode duration by not including minutes on either temporal side of the sliding windows that have low activity. To mitigate this bias, as a final step, the initial estimate was expanded/shrunk to include all adjacent minutes that show less than 25% activity so long as they were after (when available) the evening button press (indicating the subject’s intended start of attempting to sleep) and before their waking button press. A button press was considered available if the device successfully recorded a button press within sixty minutes of the estimate. The duration of the major Sleep Episode after these corrections is recorded in the data output files as the automatically generated *SleepDuration* with its beginning (*SleepOnset*) and end (*SleepOffset*) times.

Figure 3 displays examples of the initial automatic estimate of the Sleep Episode (yellow bottom bar) and the final corrected Sleep Episode (middle green bar) in the third row of Panel A, B, and C. A third estimate, shown as a blue bar, expands from the final Sleep Episode to include adjacent minutes when the activity is below the 50% threshold, such as occurs when individuals are resting in bed but not yet asleep. We store the beginning of the expanded epoch as the *BedrestOnset*, the end as the *BedrestOffset*, and the duration as the *BedrestDuration*. For some purposes, the time from the beginning of the *BedrestOnset* to the *SleepOnset* can be used as a distinct measure (e.g., *SleepOnsetLatency*). These variables are summarized in Table 1.

**Figure 3.**
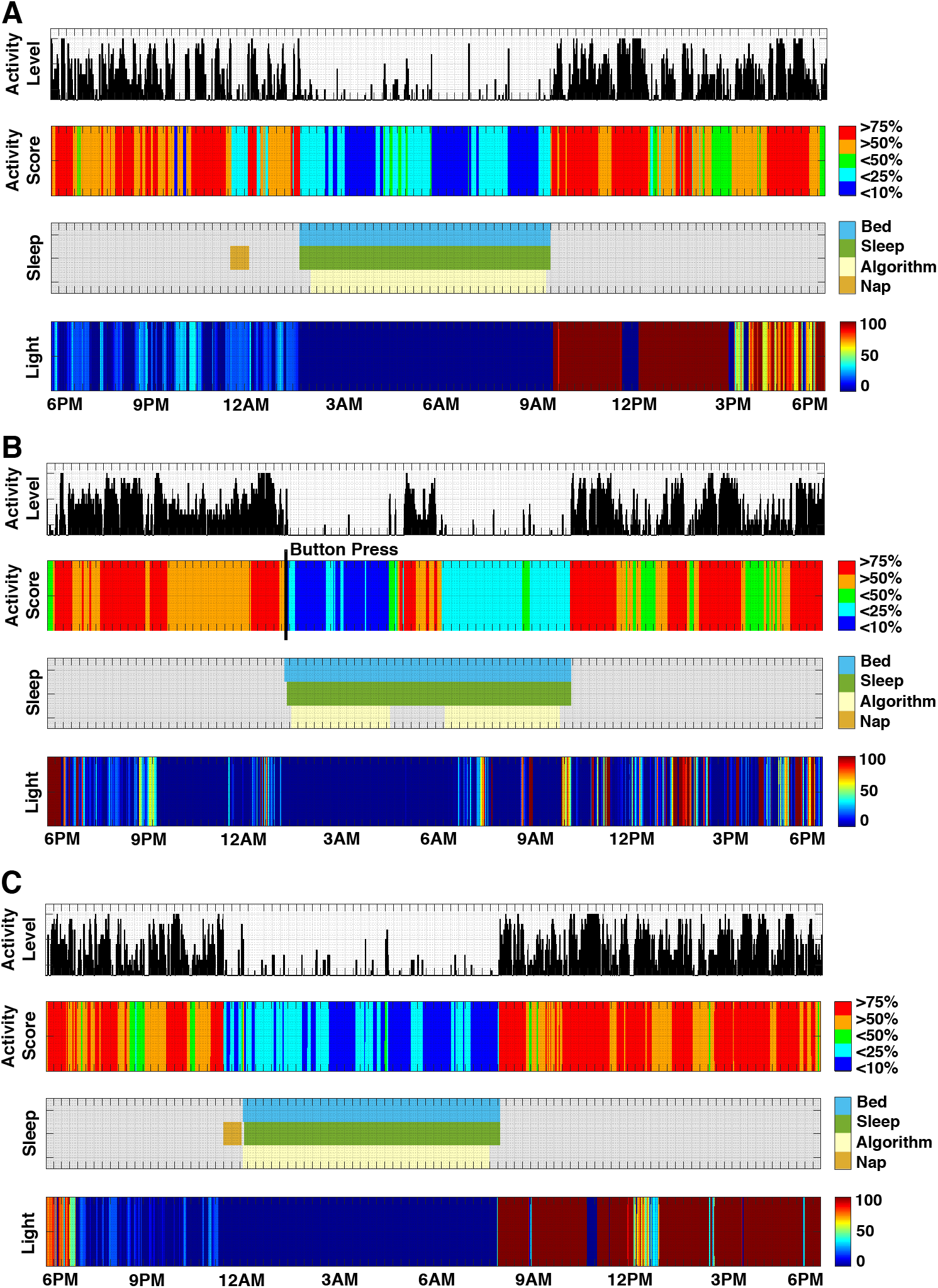
Examples of daily sleep reports. Each panel A-C displays a sleep report for a single 24-hour day of a participant. All data are from the watch. For each panel, the top row displays the activity level over time as the power of one minute (*Activity Level*). The activity score is shown in color in the 2nd row, with the low activity periods quite apparent in blue (*Activity Score)*. The third row (*Sleep*) displays the actual estimates of the Sleep Episode derived fully automatically from the activity scores and the button presses (when available). The initial provisional estimate of the Sleep Episode (called Algorithm) is displayed in yellow, with the automatically expanded Sleep Episode above it. The Sleep Episode is the main estimate used for the calculation of *SleepOnset, SleepOffset*, and *SleepDuration*. The Bedrest Episode is also shown, which extends beyond the Sleep Episode to include adjacent medium activity periods. The final row (*Light*) shows the light levels.

The estimated light exposure level from the watch is also shown in Figure 3, as well as the timestamps of the recorded button presses. The light level is not used by the algorithm to estimate the Sleep Episode but is visualized since it can aid manual adjustments that are applied during Quality Control. An assumption of the present approach is that a single Sleep Episode will occur that is usually 100 minutes or more; the limitations of this simplifying assumption will be discussed.

### Smartphone Data

In addition to wearing the actigraphy watch, most of the individuals in Studies 1 and 2 installed the Beiwe application on their smartphones. The Beiwe application was configured to passively collect phone on/off times and the GPS location of the phone. An independent pipeline was used to securely analyze the GPS data and extract the most visited places by the individual during the study and estimate their major locations every 12 min. Then a daily map was color-coded based on the presence of the individual at those points of interest. The GPS map provides the time zone of the locations where the subject has visited and evidence of location stability or movement around the time of the major Sleep Episode.

When participants traveled across time zones, a challenge arose as a matter of practice: when the new time zone is behind the old time zone, the data shifts back and overwrites the previously recorded data, and when the new time zone is ahead of the old one, the data shifts forward, and there will be a missing data gap. The same issue occurs for Standard/ Daylight Saving Time transitions. The time shifts occurring during the actual days of travel, especially when travel occurs by plane, are challenging to incorporate, and we considered these days as missing data with the days before and after being retained with data shifted to reflect the Time Zone experienced by the participant. Alternate goals, such as estimating circadian rhythms, may be better served by analyzing the data in a continuous fashion and will be different from the present focus where the discrete daily patterns require these practical adjustments.

The smartphone data from Beiwe also provided relevant data for validation, including accelerometer and phone use data (via the recorded lock-unlock events). The DPSleep pipeline allows the integration and visualization of smartphone data when available. To integrate accelerometer data for the present paper’s validation purposes, a simple standard deviation analysis was applied to the phone acceleration along the x-axis as a representative of the acceleration score to recognize minutes of high movement. The distribution of acceleration score in all minutes during the study for each individual was used to find minutes with greater than 75 and 90 percentile movement (normalized to the individual). These minutes were color-coded in yellow and red, respectively, contrasting the lower acceleration minutes in gray, and plotted in relation to the daily Sleep Episode estimates (Figure 4). Additionally, the locked-unlocked events of the phone document when the phone is in use. Phone in-use time was defined as the time between every consecutive unlocked-locked events, lasting no more than 15 minutes. This is to soften the strong assumption of the phone being in-use all the time after the unlocked event and before it is locked again. The daily map is then color-coded to show the locked-unlocked minutes, in red and blue, respectively, in addition to in-use minutes in green (Figure 4). These data are used in the present paper to build confidence in the DPSleep estimates of the major Sleep Episode. They may also have utility for understanding the relation of digital technology use to sleep patterns, for example, as might occur if individuals use their phone sporadically at night.

**Figure 4.**
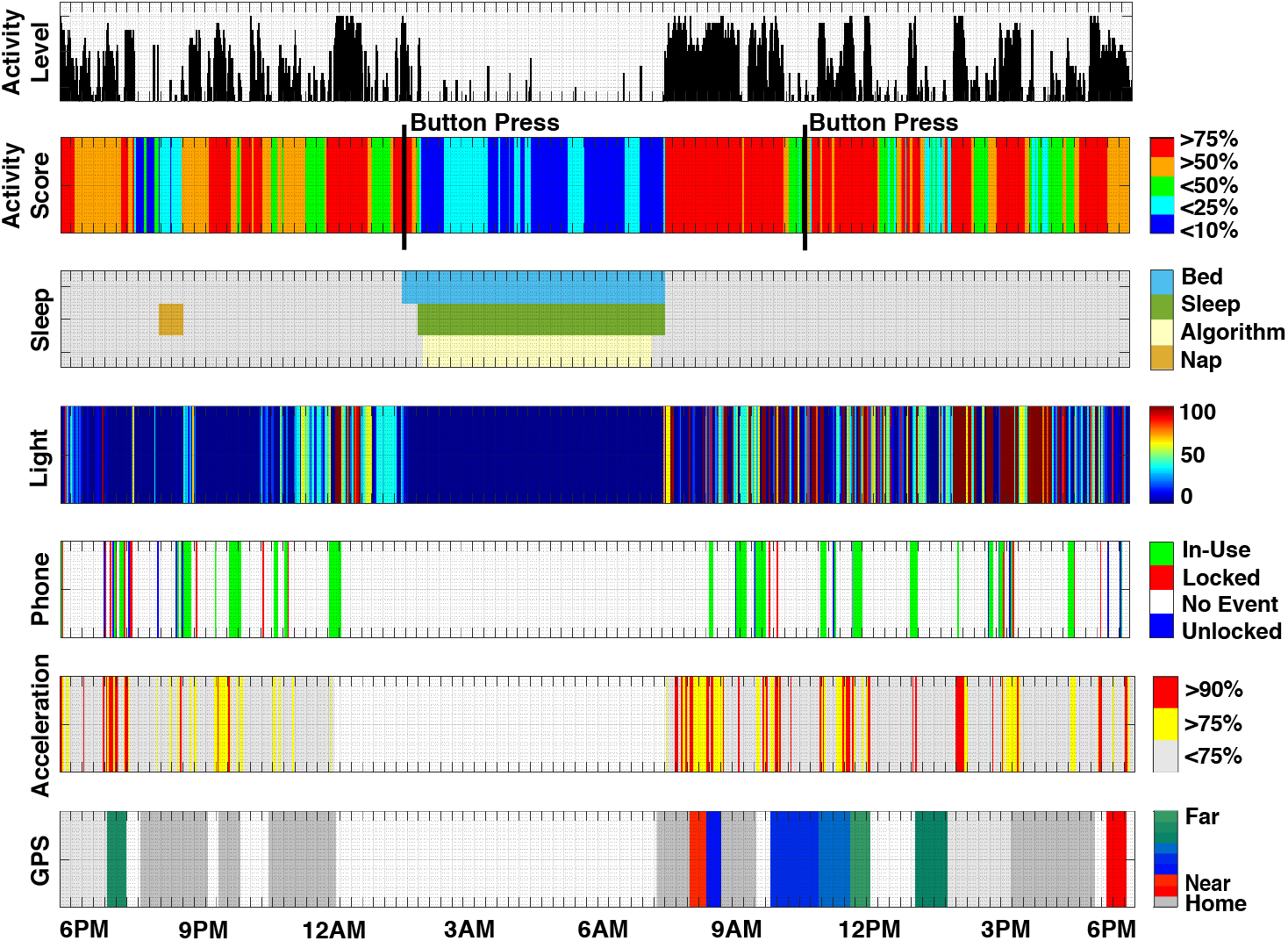
Example of a daily sleep report that includes phone use and GPS data. Data for a single 24-hour are displayed similar to Figure 3 with additional data obtained from the smartphone of the individual. The first four rows are identical to Figure 3. The fifth row (*Phone*) displays lock-unlock phone events. The sixth row (*Acceleration*) displays data derived from the accelerometers within the phone. The final row (*GPS*) shows the clustered frequently visited locations of the individual relative to their estimated home location (dark gray), with different colors showing different distances from home. Missing GPS data are shown in white and light gray indicates the available coordinates that are not among the frequently visited locations by the individual.

### Sleep Estimation Quality Control

DPSleep should not be expected to deliver perfectly accurate output when operating in the automatic mode. Several assumptions are made, and the structure of an individual sleep night can be complex. Instead, DPSleep provides an elaborative day-by-day report, examples of which appear in Figure 3. The user can decide about the confidence of the estimated Sleep Episode and revise the results manually if necessary. DPSleep includes editing tools. Every page of the report presents the data about one day from 6PM of the previous day to 6PM of the original day. The report can also include, when available, smartphone data, as illustrated (Figure 4). The daily report is a significant help to the investigators to decide about, and increase the precision of the sleep estimation results based on the availability and richness of the data.

All results in this paper have been quality-controlled by two individuals independently relying on only the watch actigraphy data and not any ancillary data from the smartphone to make modifications. Thus, the data are analyzed here as would be from any typical study that only obtained watch actigraphy data. The guidelines we used for our quality control are described in Appendix II. Figure 5 illustrates plots of the sleep duration across nights before and after manual adjustments for each of six individuals in Study 1. As shown, most nights show identical values before and after QC, meaning no adjustment was required, while many others show slight adjustments. For several nights, a large adjustment was required (e.g., in Figure 5 S6, there is an outlier value; in Figure 5 S5, there are several values that were substantially corrected). The automated values showed a correlation with the final corrected values that ranged between r = 0.92 to r = 0.98. Figure 6 illustrates examples of errors encountered that required manual adjustments. Manual adjustments were made for 18% of the nights. As is illustrated in Figure 5, while approximately one in five nights were adjusted, most were small adjustments that would not impact most analyses. About 9% of the nights were adjusted to change the major Sleep Episode estimate by greater than 1 hour.

**Figure 5.**
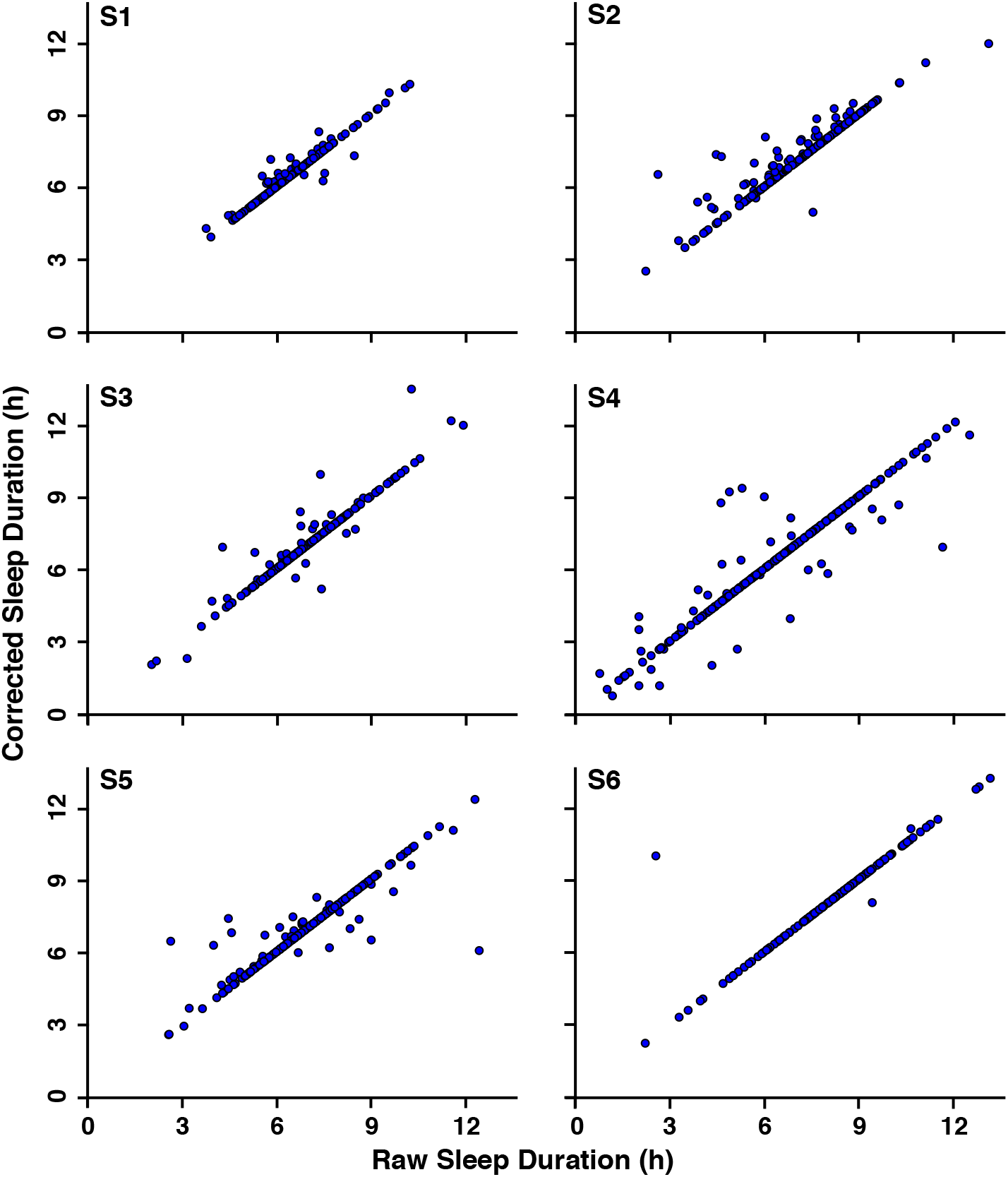
Effects of manual quality control adjustment on Sleep Duration. The Raw Sleep Duration estimates for each of the six subjects of Study 1 are plotted against the Corrected Sleep Duration estimates after quality control adjustment.

**Figure 6.**
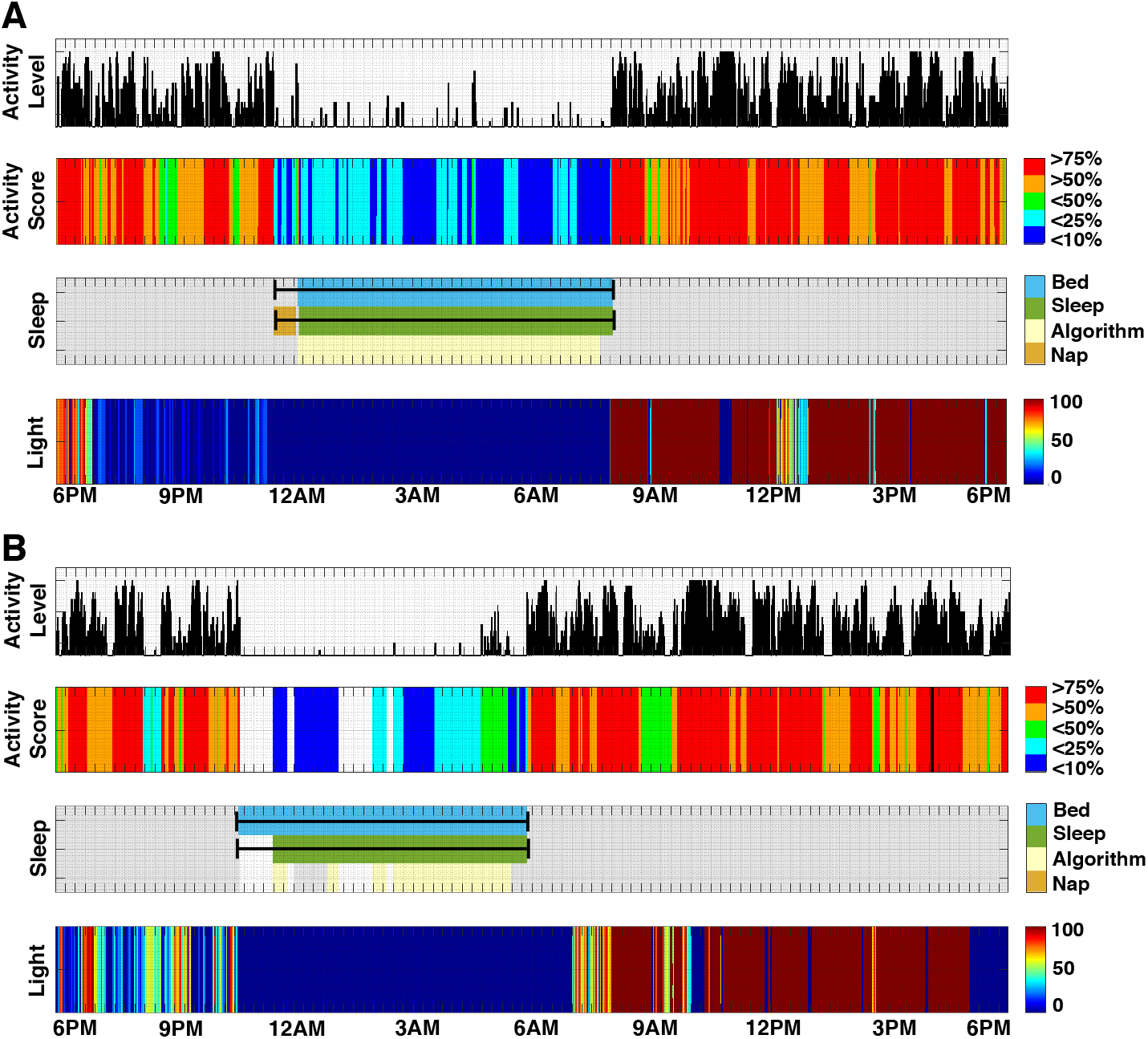
Examples of manual adjustment of the Sleep Episode. Two examples of adjustments to the Sleep Episode are illustrated. The adjusted episode is noted by the black line with marked ends plotted on top of the automatically generated Sleep Episode (green bar) and Bedrest Episode (blue bar). (A) A case where the automated process identified a short sleep adjacent to the main Sleep Episode. The two were concatenated into one extended Sleep Episode. (B) A case where the stillness during sleep appears to fall below the threshold and gets erroneously labeled as a wrist-off event (white, missing data). However, the intermittent movements suggest the watch is being worn. Manual adjustments only used the data from the watch actigraphy device.

### Data Security

Throughout the entire analysis, the pipeline is designed to securely handle the data without exposing any identifiable information. Days are reported as the days of the study relative to the consent day of the individual. We consider the GPS data identifiable not only when the actual coordinates are presented but also patterns of the daily maps as presented with significant (for the individual) locations. Therefore, the pipeline was designed to work with the encrypted data and avoid saving any of the coordinates or maps as unencrypted scratch files. Final results are saved directly to an encrypted format.

### Within-Individual Statistical Modeling

To analyze the longitudinal association between the actigraphy-based sleep estimation and the self-report data, a simple individual-level linear model was used with the self-report sleep quality as a predictor and actigraphy-based sleep duration as the outcome. This simple linear model was selected after using mixed linear model analysis to account for the time and auto-correlation of the observations. Categorical day of the week was included as a covariate to account for weekly structure related to course schedules and weekday-weekend differences. Only data collected during the academic year were included (i.e, fall and spring semesters, including exams periods and Thanksgiving and spring breaks, but excluding winter and summer breaks) to account for differences between the school year and extended school breaks. All the analyses were conducted in R using the ‘stats’ package *lm()* function (R Core Team, 2019; version 3.6.1).

## Results

### Longitudinal Activity-based Sleep Estimates Within Individuals

The longitudinal sleep patterns and daily sleep maps are the key outcomes of the DPSleep pipeline. To show the performance of the pipeline, four examples of processed data are shown for individuals from Study 1 (Figures 7, 9, 11, 13). The data are from the full academic year with travel to and from campus and across Daylight Saving Time. Each figure contains three plots. The first is the color-coded daily map of activity scores that show the minutes with high (>50% orange, >75% red), low (<25% cyan; <10% blue), and medium (25-50% green) activity. The second plot is the raw estimated Sleep Episodes for each day. The final plot is the Time Zone adjusted and quality-control adjusted longitudinal plot of sleep behavior. This final plot represents the data used for all quantitative analyses and validation.

**Figure 7.**
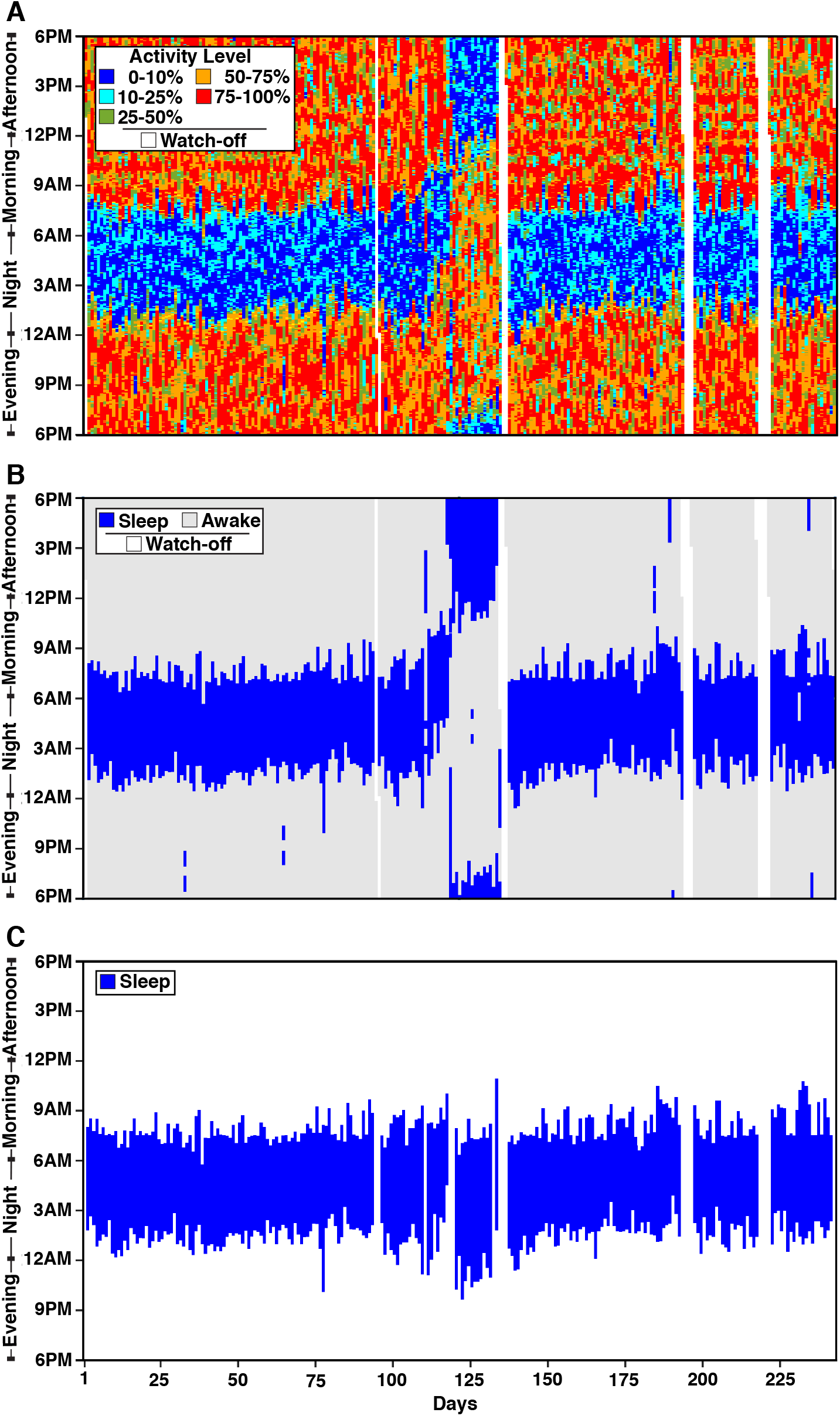
Longitudinal Sleep Episode estimates in S1. Three panels display longitudinal Activity Score and Sleep Episode estimates for 243 days. Day 1 is near to the 9^th^ day after the beginning of the semester. Winter break falls near days 102 to 133. Each vertical line displays data for a 24-hour day. (A) The top panel displays the continuous Activity Scores colored by the threshold for each day. For each day, the plot begins at 6PM at the bottom and ends at 6PM, allowing the nighttime sleep period to plot in the center of the graph. Over days the low activity band (in blue) is relatively consistent except for a dramatic shift at Day 118, which is attributed to travel outside of the Time Zone. (B) The middle panel displays the same data revealing the automated detection of Sleep Episodes and Nap Periods. (C) The bottom panel C displays the Time Zone adjusted and fully quality-controlled estimates of the final Sleep Episodes.

**Figure 8.**
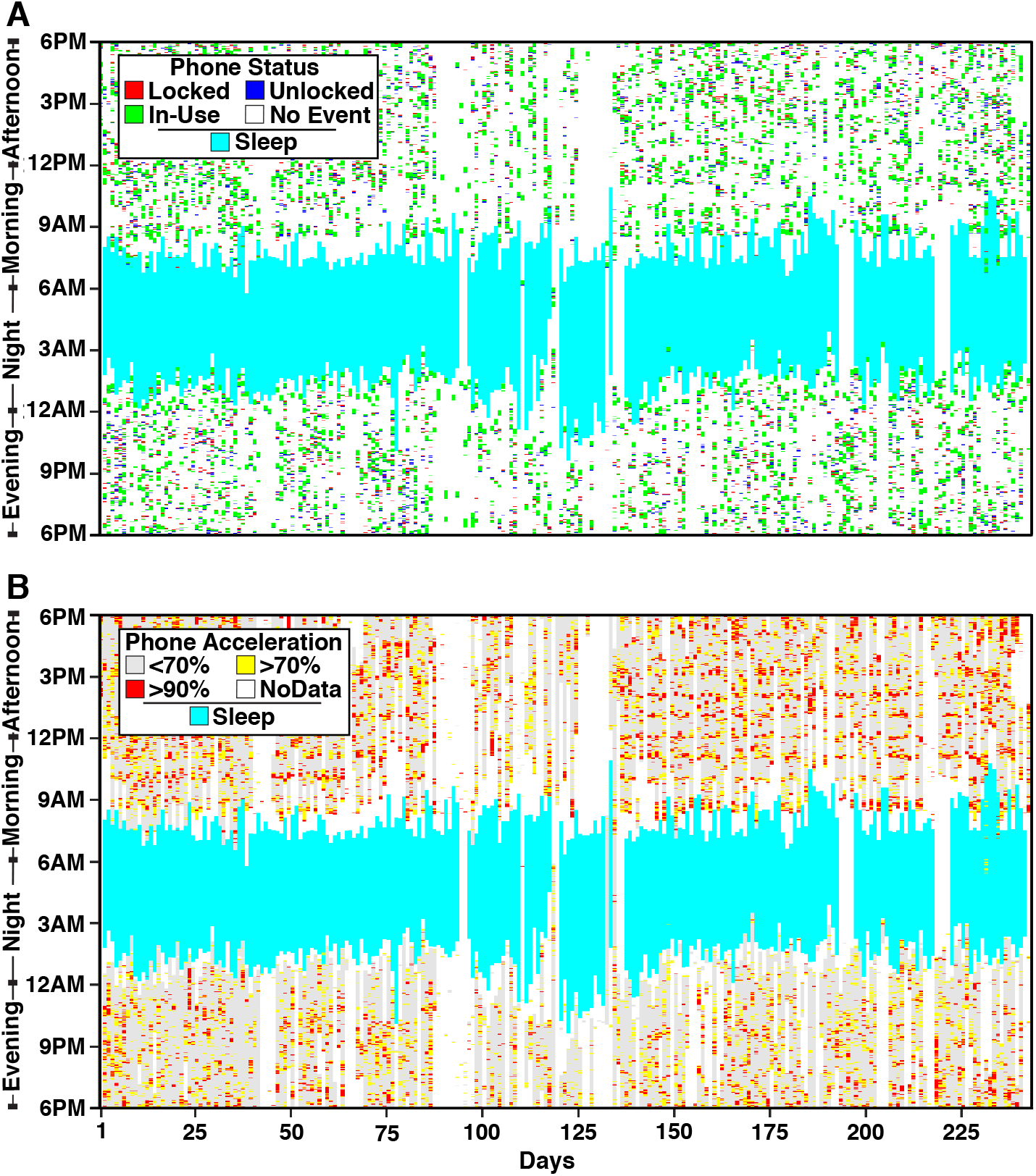
Longitudinal Sleep Episode estimates in S1 in relation to phone use. The Sleep Episodes from Figure 7C are plotted (light blue) in relation to independently estimated phone events. (A) The Phone Status is shown with colored hashing for when the phone is Locked (red), Unlocked (blue), and In-Use (green). Note that there is virtually no phone use during the estimated Sleep Episodes. The phone use events, on many nights, occur up until and just before the beginning of the Sleep Episode. However, on most mornings, there is a gap between when the Sleep Episode ends, and phone use begins, which often begins abruptly near to 9AM, possibly reflecting phone use begins with an alarm-triggered event. (B) Phone Acceleration data are plotted and also reveal virtually no phone movement during the Sleep Episodes

**Figure 9.**
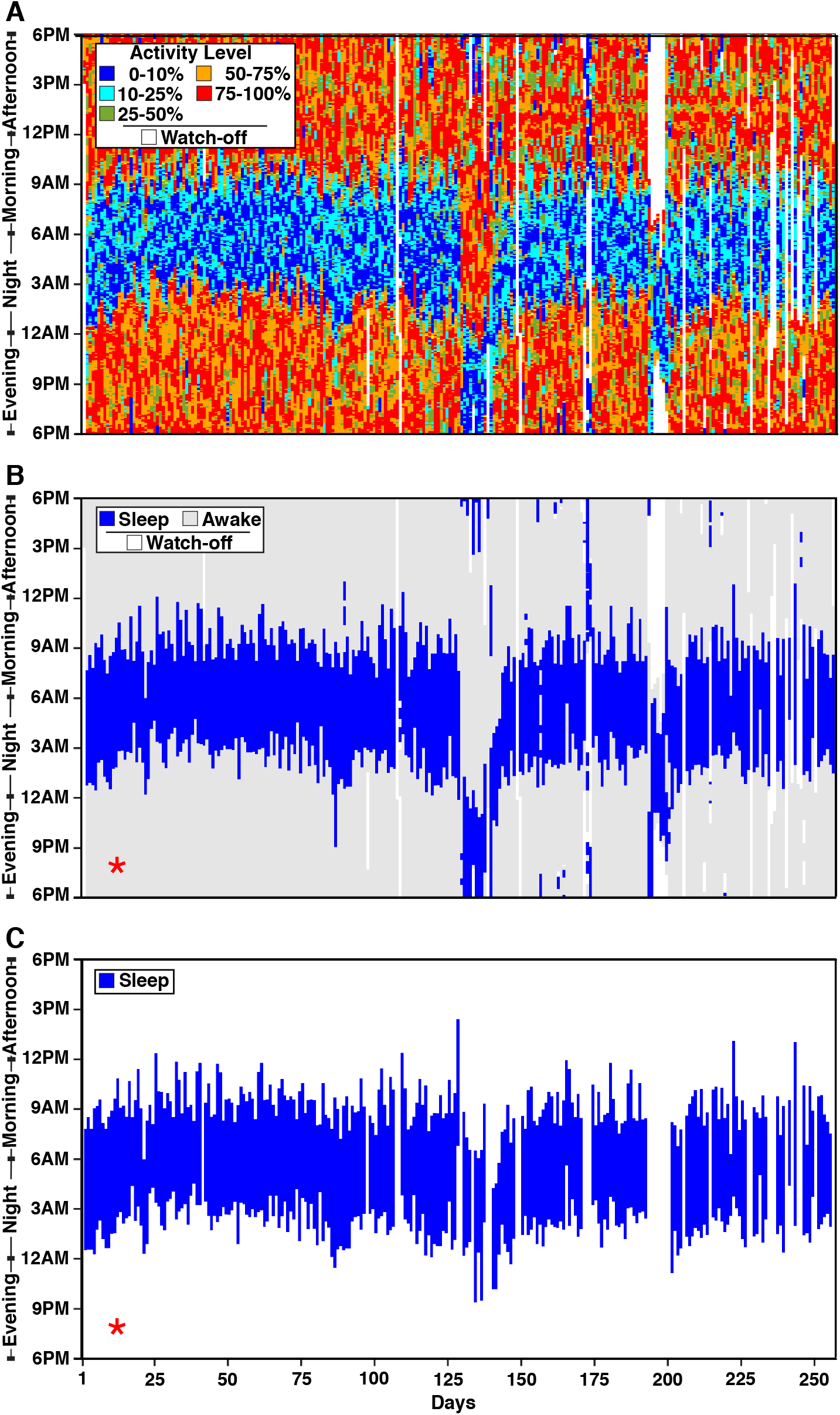
Longitudinal Sleep Episode estimates in S2. Longitudinal Activity Score and Sleep Episode estimates for 257 days are plotted in the same format as in Figure 7. Day 2 is near to the beginning of the semester. Winter break falls near days 112 to 143. An interesting feature in this individual is the extended wrist-off periods from Days 194 to 201. As can be seen in panel C, these days in their entirety are excluded from the final analysis. Another interesting feature in this individual is the gradual shift to a later Sleep Episode from Day 1 to about Day 25, indicated with the red asterisk in B and C. The later Sleep Episode is then maintained for much of the semester.

**Figure 10.**
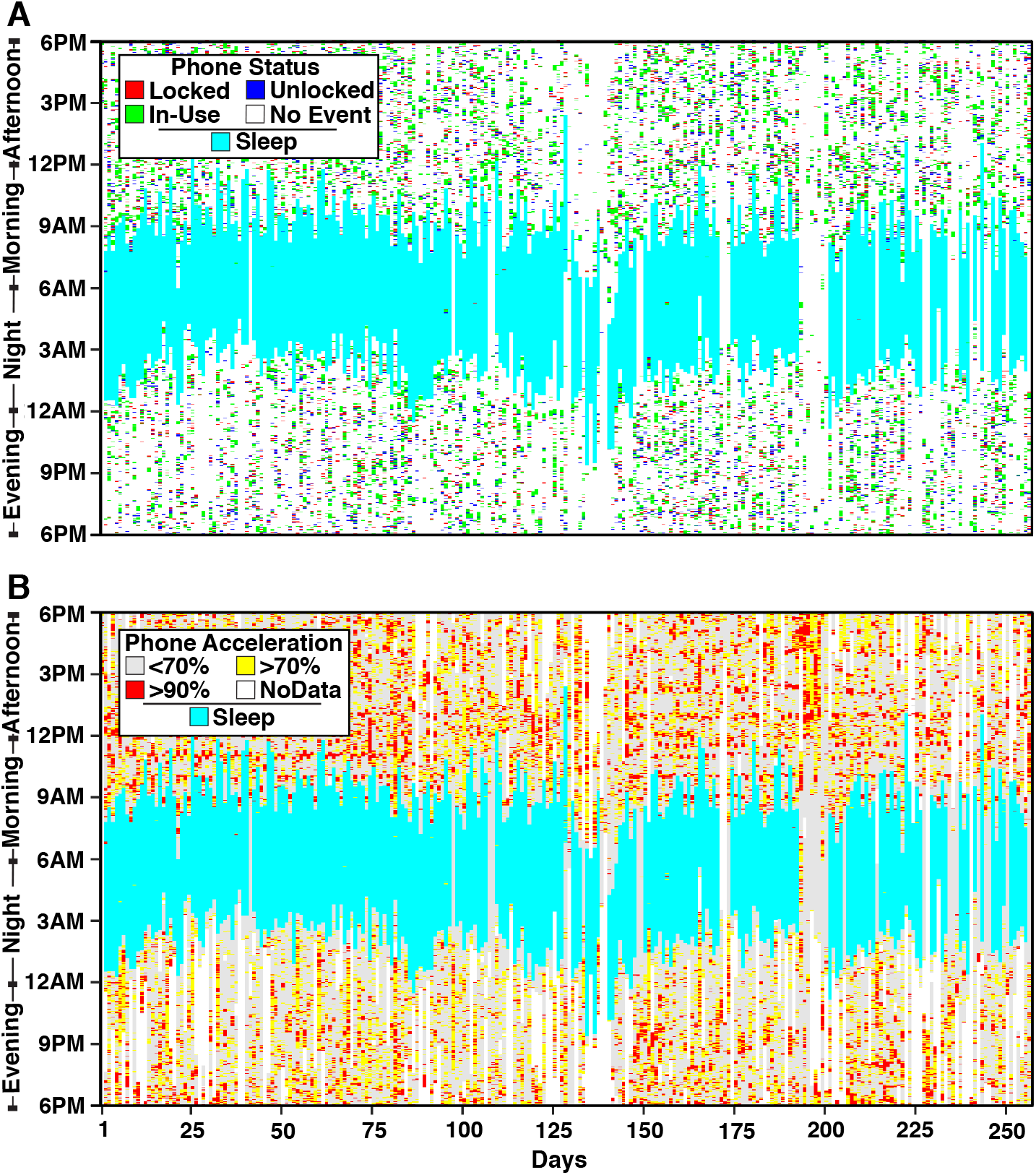
Longitudinal Sleep Episode estimates in S2 in relation to phone use. Phone use data are plotted in the same format as Figure 8. Again, there is minimal phone use during the Sleep Episodes, consistent with this individual not using their phone during the nighttime. In contrast to S1, Phone Status events begin almost immediately following waking, suggesting that the individual starts using her or his phone as soon as sleep ends.

**Figure 11.**
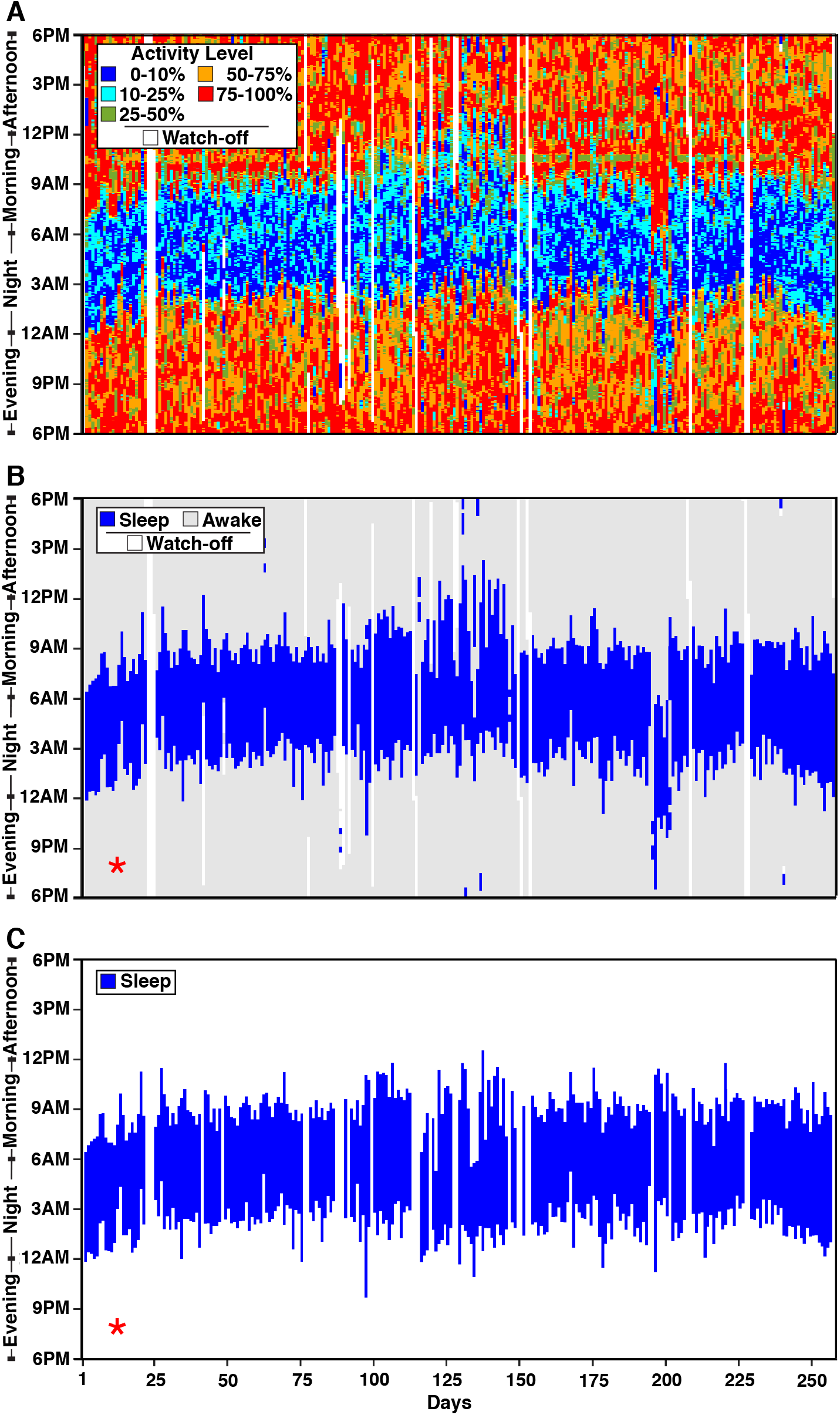
Longitudinal Sleep Episode estimates in S3. Longitudinal Activity Score and Sleep Episode estimates for 258 days are plotted in the same format as in Figure 7. Day 4 is near to the beginning of the semester. Winter break falls near days 114 to 146. Similar to S2, this individual shows a gradual shift to a later Sleep Episode from Day 1 to about Day 25, indicated with the red asterisk in B and C.

**Figure 12.**
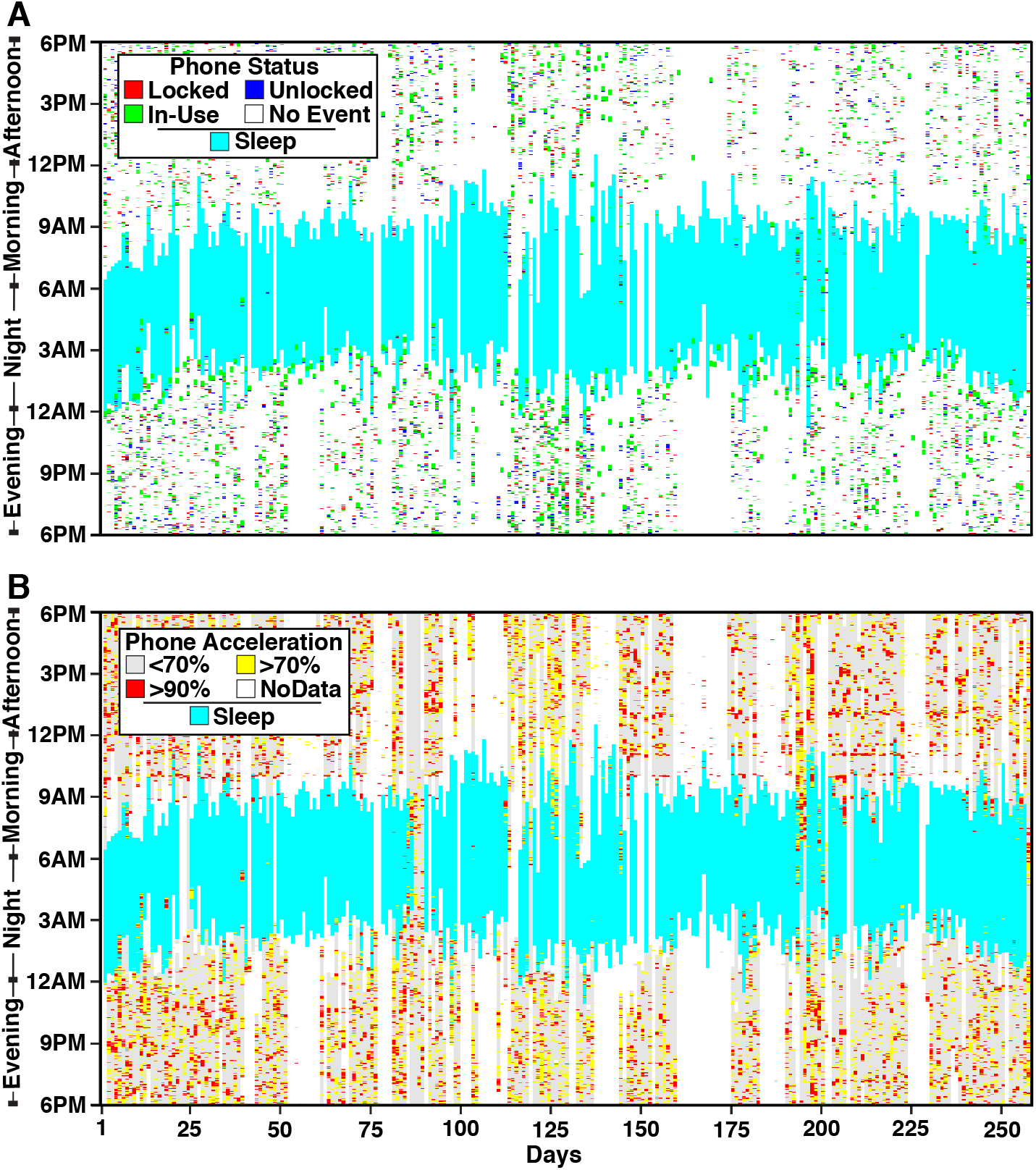
Longitudinal Sleep Episode estimates in S3 in relation to phone use. Phone use data are plotted in the same format as Figure 8. In this individual, phone use often ends well before the estimated Sleep Episode begins. In addition, phone use shows a structured pattern on wakening with often-recorded acceleration events at 9AM, then a gap, and then another bout of events at 10AM.

**Figure 13.**
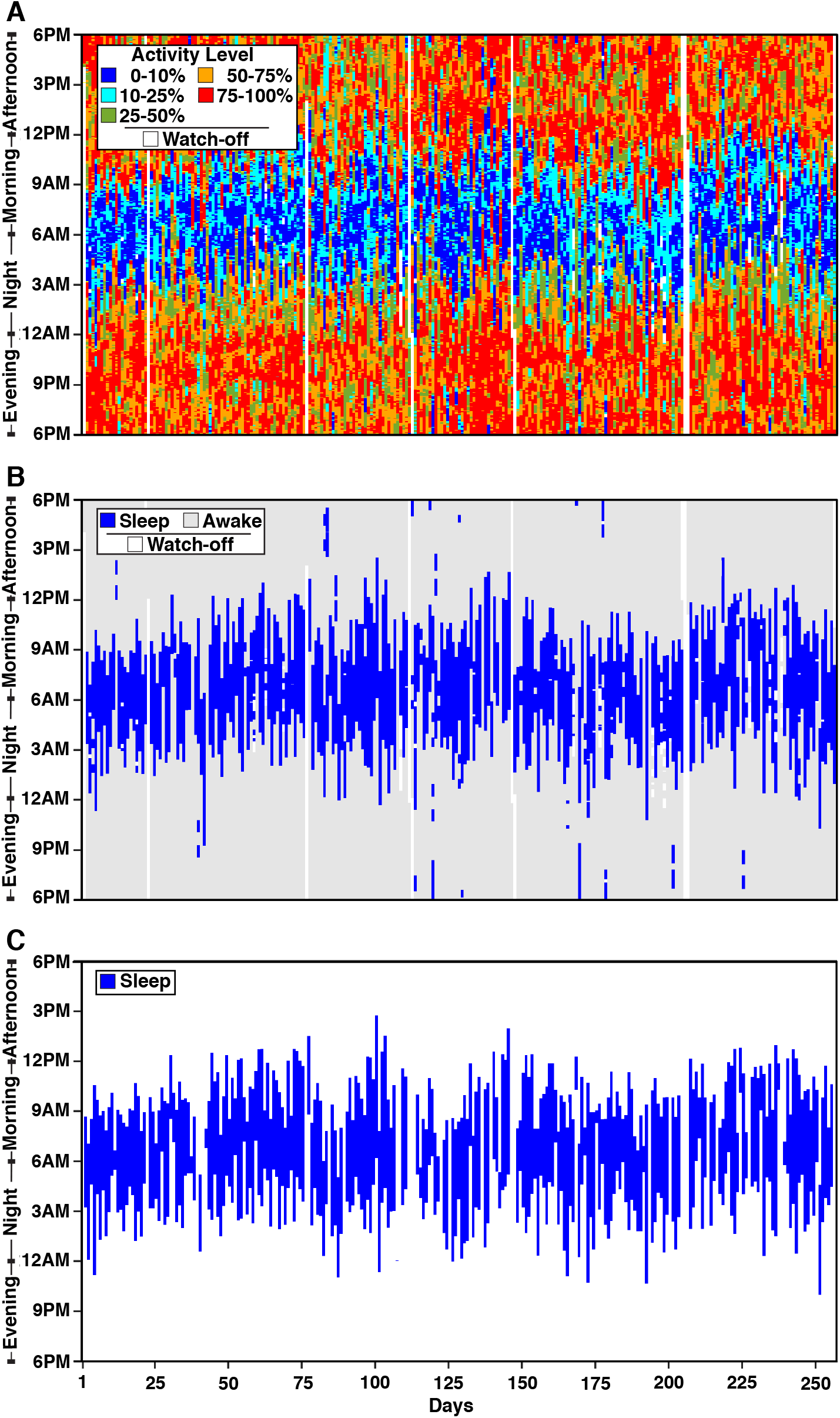
Longitudinal Sleep Episode estimates in S4. Longitudinal Activity Score and Sleep Episode estimates for 257 days are plotted in the same format as in Figure 7. Day 2 is near to the beginning of the semester. Winter break falls near days 112 to 143. This individual shows highly irregular Sleep Episodes.

Figure 7 shows the sleep data for S1 with a highly regular sleep pattern where the subject goes to bed sometime between 1AM and 2AM and wakes up consistently near 8AM. Every 6-7 days, the subject wakes up about an hour later than usual, which could be the indicator of the weekends. Figure 9 shows the sleep data for S2 with a less regular sleep pattern where, during the semester, the subject goes to bed between midnight to 4AM and wakes up between 9AM and noon. The effect of beginning the academic year is notable with a gradual transition to a later sleep time (first 25 days). The break and travel also affect the sleep schedule significantly during days 125-150. The subject does not wear the wristband during the day for the travel days of 190-200, which are coded as missing data. Figure 11 shows the sleep data for S3, whose sleep schedule is comparably influenced by the academic calendar. Figure 13 shows the sleep data for S4, who displays the most irregular sleep pattern. There are some nights when this subject goes to bed at around midnight and does not wake up before 9AM, and other nights when the individual does not go to sleep before 6AM and sleeps only a few hours. The irregular sleep pattern does not appear to be related to the academic calendar since the subject displays a similar sleep pattern during breaks.

### Smartphone Use Tracks Sleep Episodes

One way of validating the estimated Sleep Episode arises from the independent phone data collected simultaneously through the Beiwe application on each individual’s cell phone. Even though these data do not continuously measure activity, since the phone can be put down, they do indicate the minutes during which the individual is clearly not sleeping. Figures 8, 10, 12, and 14 plot the smartphone data for the four subjects analyzed above. In each plot, the upper panel shows the Phone Status via the locked-unlocked events. When the phone is not used, there should be no changes in the Phone Status. The lower panel displays Phone Acceleration that, like the wristwatch, provides a measure of dynamic movement when the phone is picked up, used, or is moving with the subject’s body. The missing signal instead of the low signal on the acceleration maps could be due to the subjects turning off their phones, or because when the phone is not in use or does not move for a certain time period, the operating system can enter a power-saving mode that temporarily disables data collection from sensors.

**Figure 14.**
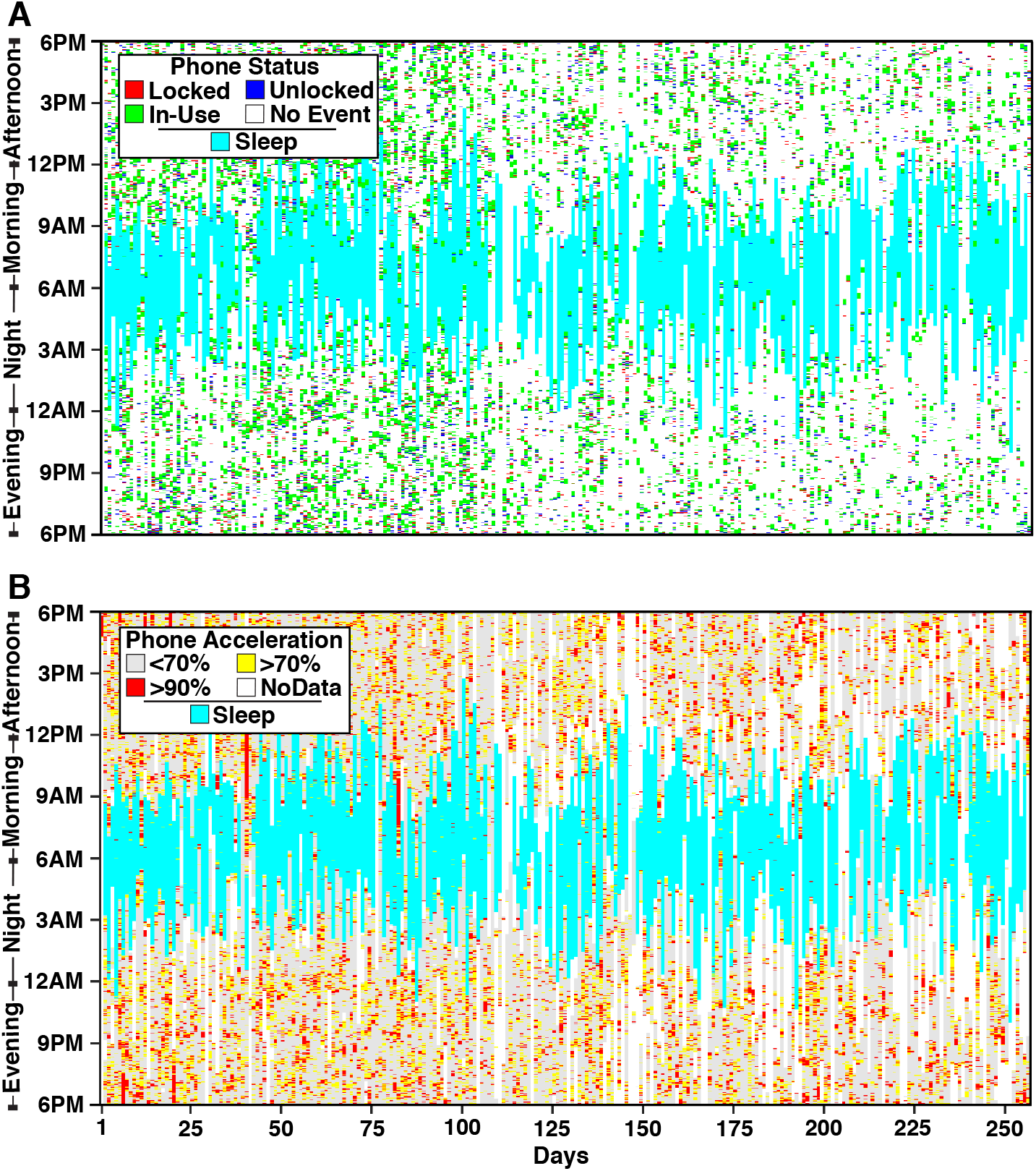
Longitudinal Sleep Episode estimates in S4 in relation to phone use. Phone use data are plotted in the same format as Figure 8. Note that the Phone Status (A) and Phone Acceleration (B) events track the irregular Sleep Episodes.

In all subjects, the phone activity measures were generally outside the time of the estimated Sleep Episodes and tracked the variations in sleep period from night to night. S2 and S3 (Figures 10 and 12) show this quite clearly since their sleep patterns change gradually during the first 25 nights of measurement. The phone activity measures track these transitions. S4, who showed the most erratic sleep patterns, also demonstrated quite clear evidence that phone use was frequent and intensive only outside the time of the major Sleep Episodes (Figure 14). Beyond the general correspondence between phone and sleep, there were also interesting features in the details of the phone use that are relevant for comparing the activity- and phone-based data types.

One detected feature is that the subjects occasionally used their phones for brief periods during the major Sleep Episode. A good example of this is S3 who showed short periods of phone use on occasion during the middle of her or his Sleep Episode (Figure 12; Days 19, 26, and 40). Another detected feature is that phone use does not precisely align with the beginning and end of the Sleep Episode. We suspect that this is a genuine difference in phone use patterns between subjects rather than a technical error. For S2, phone use tracked the major Sleep Episode remarkably well with phone use events occurring until the Sleep Episode began and beginning shortly after the Sleep Episode ended (i.e., awakening) (Figure 10). This particular individual also showed virtually no phone use during the Sleep Episode, yielding an extremely tight coupling of the two independent streams of data. However, other individuals showed systematic differences. For example, S1 (Figure 8) showed a gap between the estimated wake time and the first evidence of phone use. Moreover, phone use across days showed a patterned start at 9AM for most days, even when the Sleep Episode ended earlier. This individual likely does not start using their phone immediately upon waking; there may be a phone alarm set at 9AM leading to a consistent first use at the same time across days. These systematic differences show that combining the data can reveal the interaction between device use and sleep pattern (e.g., by detecting the tendency to briefly use the phone during nighttime wakening).

As an additional visualization, Figure 15 shows data from S1 with the horizontal axis representing clock time (48 consecutive hours, with the 2^nd^ 24 hours on each horizontal line repeated in the 1^st^ 24 hours of the next horizontal line) and day number down the y-axis; this is the plotting convention often used by investigators interested in circadian rhythms (Dagan and Borodkin, 2005). DPSleep allows plotting using either lateral (as in most figures in this paper) or horizontal (e.g., Figure 15) conventions.

**Figure 15.**
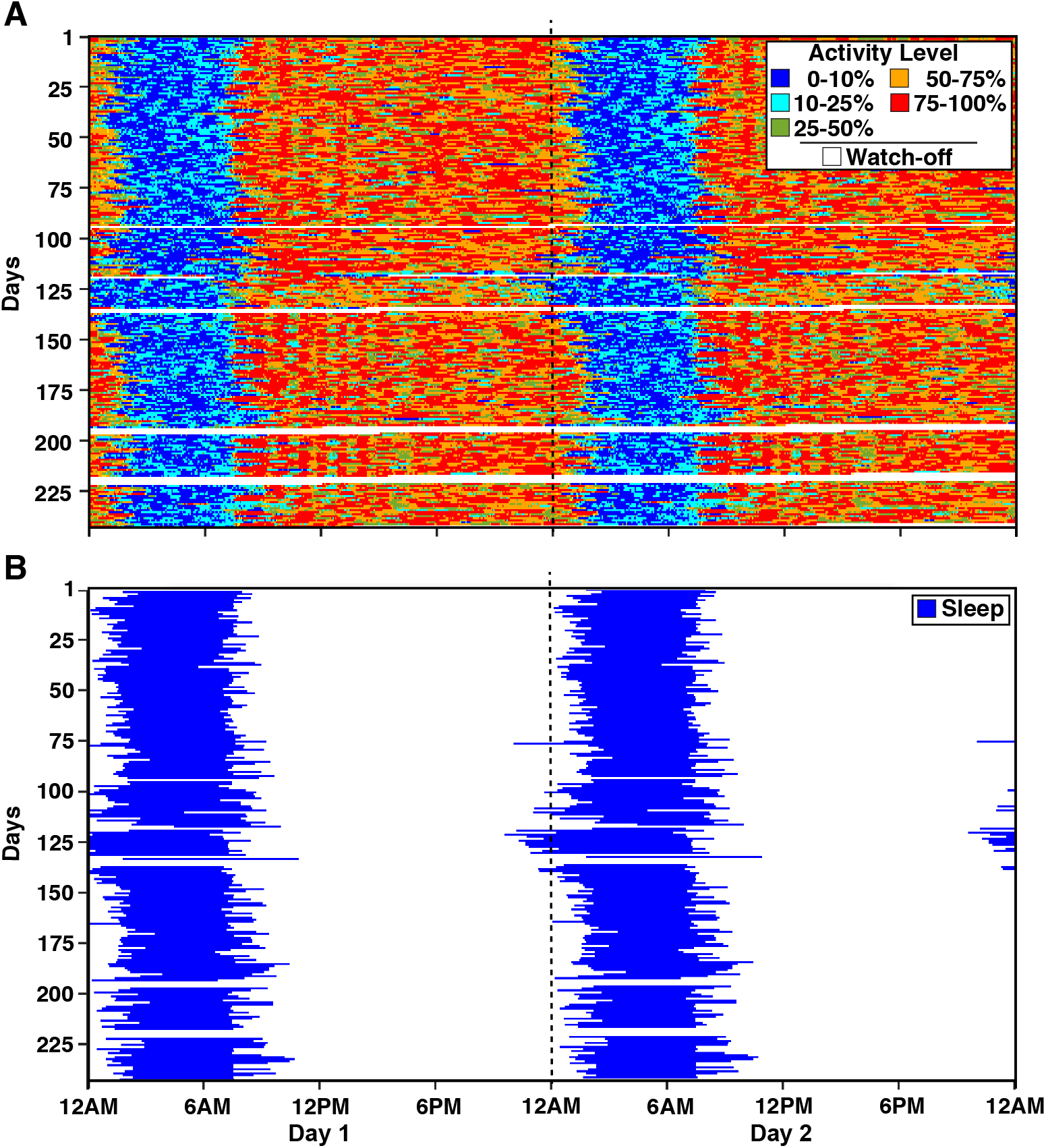
Two-Day column formatted Longitudinal Activity Score and Sleep Episodes. Color-coded Activity Score (A) and the estimated Sleep Episode (B) are presented in a format that is standard in the sleep medicine community. In this format, the days of the study are presented as the rows from top to bottom and every row contains the data of two consecutive days; the first column is the original day and the second column is the next day both presented from 12AM to 12AM. The format allows the investigators to focus on the study nights with the vertical blue band in the middle of the plot, without missing any information before and after the night. This individual (S1) Sleep Episodes are shifted to the right of the center because their sleep onset time is typically after 1AM.

### Actigraphy Measures of Sleep Duration Track Self-Report Sleep Quality

In each individual, the DPSleep pipeline yielded an estimate of the Sleep Duration (*SleepDuration* in Table 1), the time difference between the *SleepOnset* and the *SleepOffset*. Each day, the individuals also reported the quality of the previous night’s sleep on a Likert Scale (Appendix I). Figure 16 shows the variation in Sleep Duration across days in each of the six individuals from Study 1 as well as their self-report Sleep Rating. Missing data excluded from the analysis include the data from winter break (the gaps near Day 120) and non-compliant data (e.g., the wristband was removed or the survey was submitted the following day).

**Figure 16.**
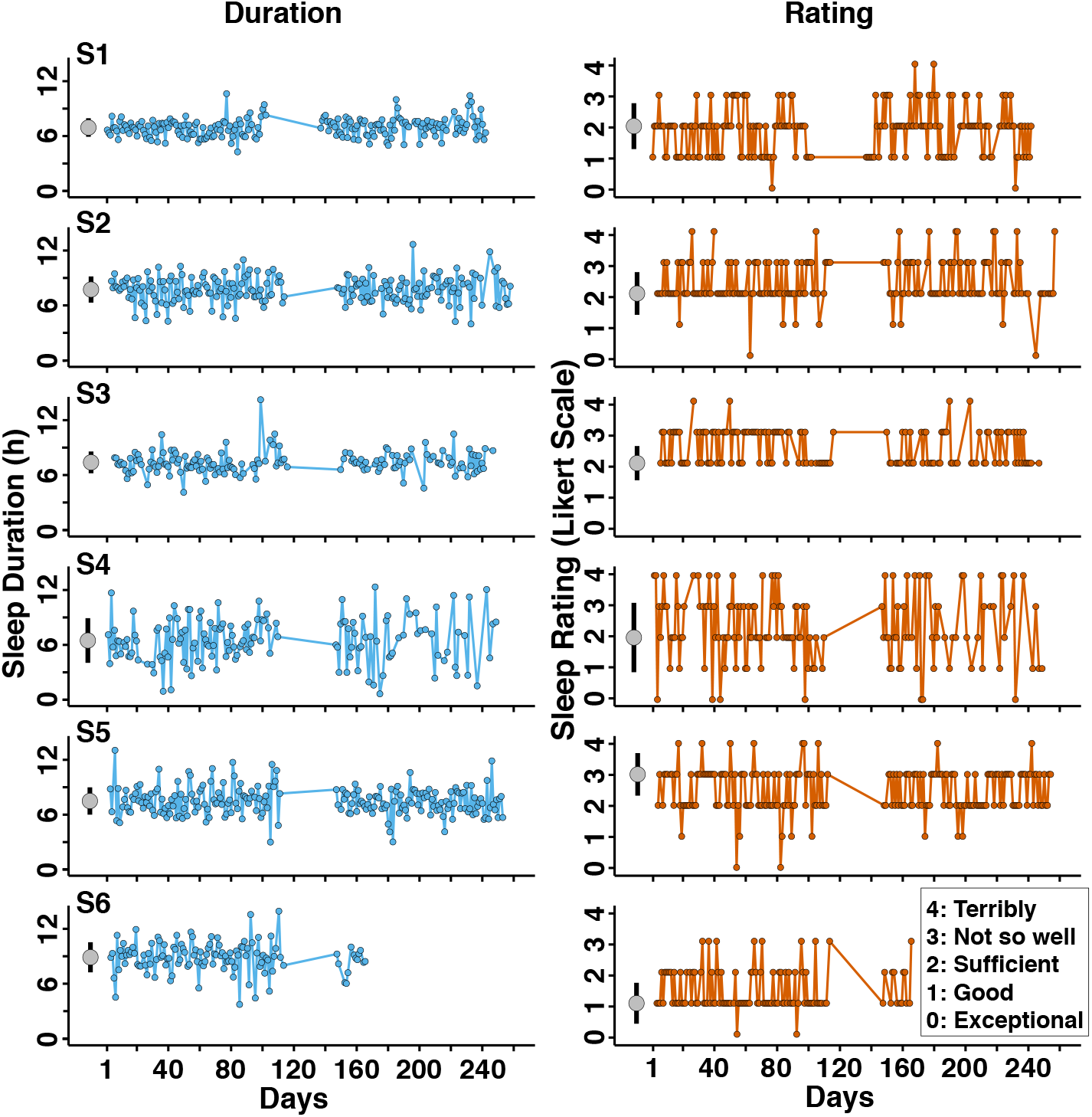
Longitudinal Behavior of Sleep Duration and Sleep Rating. The time course of each subject’s objective Sleep Duration (in minutes) as measured by actigraphy is plotted (left column) next to the same subject’s self-report Sleep Rating (right column; 5-point Likert scale; 0=Exceptional, 4=Terribly). Data from winter break and non-compliant data (e.g., the watch was removed, or the survey was submitted the following day) are excluded; hence there are gaps in the data, including a large gap around day 120 (winter break). The gray circles to the left of each time course show the mean (± SD) Sleep Duration and most frequent (mode ± SD) Sleep Rating for each individual.

Individual-level linear model analysis is used to evaluate the longitudinal association between the self-report sleep quality (for both concurrent night and nights preceding) as the predictor and the actigraphy-based Sleep Duration as the outcome. There is a significant correlation between the concurrent night’s self-report Sleep Rating and the actigraphy-measured Sleep Duration in each of the six subjects (Figure 17): short Sleep Durations predict self-report ratings of poor sleep. The observation that the Sleep Rating and Sleep Duration measures track one another in healthy young individuals provides evidence of validity. No correlation or a weakly positive correlation is observed for the preceding night.

**Figure 17.**
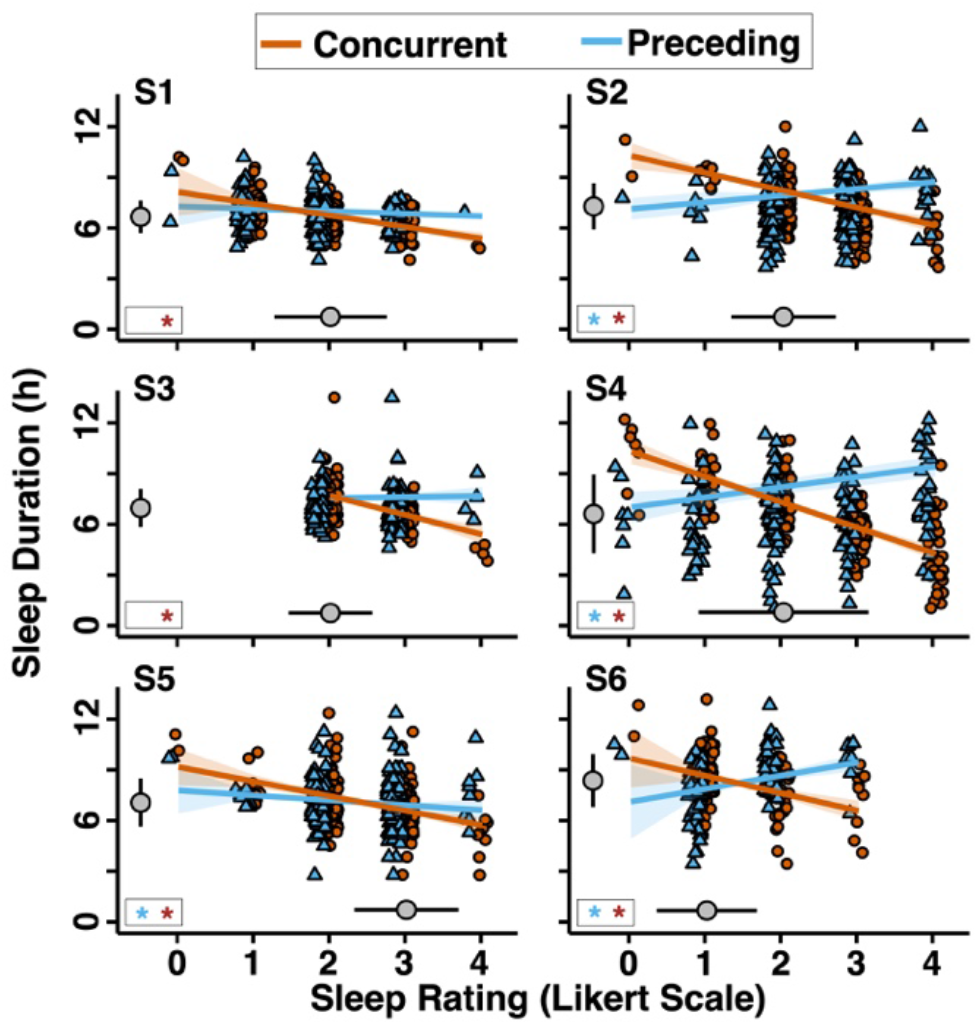
Self-report Sleep Rating predicts Actigraphy-based Sleep Duration. Individual-level linear model applied to each subject shows the association between self-report Sleep Rating and actigraphy-based Sleep Duration. In each case, the self-report Sleep Rating negatively predicted Sleep Duration when the Sleep Rating targeted the night of sleep (red line and circles). When the Sleep Rating was shifted to the next day so that the rating no longer matched the night of Sleep Duration, the relationship between Sleep Rating and Sleep Duration in most subjects showed either no relation (S1, S3) or a small positive relation, perhaps a form of sleep rebound (S2, S4, S6) (blue triangles and lines). S5 showed negative association for both nights. The larger gray circles to the left of each plot show the average (mean ± SD) Sleep Duration across the study for each subject. The larger gray circles below each plot show the most frequent (mode ± SD) Sleep Rating across the study for each subject. The bottom left box in each panel indicates which associations are significant (p<0.05).

### Sleep Duration and Sleep Ratings Show Structured Weekly Variation

One of the outcomes of accurate estimation of the major Sleep Episode in individuals is the ability to investigate non-stationary effects that fluctuate with the weekly schedule, holidays, and academic calendar (especially since subjects in Study 1 are undergraduate students). To evaluate weekly variation that might be relevant to data modeling as well as discovery insight, an autocorrelation analysis was performed. Actigraphy-based Sleep Duration shows almost no lag effect such that one night’s duration shows little association with the next (Figure 18). Interestingly, there is a notable lag effect at day 7 and day 14, suggesting a strong autocorrelation between the same days of the week. Self-report Sleep Rating shows similar lag day 7 and lag day 14 effects and also a temporal autocorrelation at lag days 1 and 2 suggesting a poor or good Sleep Rating predicted a similar rating on subsequent nights despite no evidence of autocorrelation in the actigraphy-based Sleep Duration.

**Figure 18.**
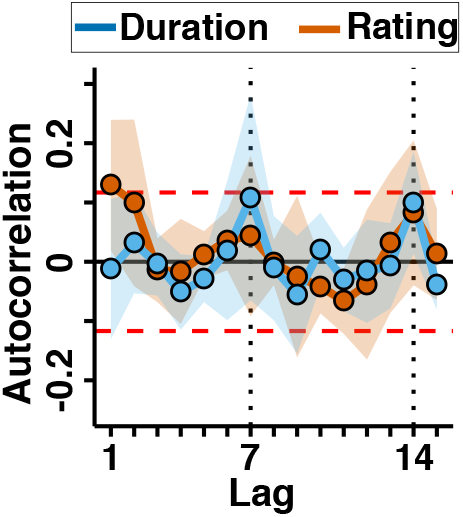
Autocorrelation (lag) analysis of Sleep Duration and Sleep Rating reveal weekday patterns. The autocorrelation, or the correlation of each variable’s time course with itself at varying lags, is plotted for actigraphy-based Sleep Duration (blue) and self-report Sleep Rating (red) averaged across the six subjects of Study 1. Shading illustrates the standard error of the mean. Increased autocorrelation values at lags of 7 and 14 days indicate that Sleep Duration and Sleep Rating for a given weekly night (e.g., Friday, Saturday) are more similar to the same weekly nights on different weeks than to the immediately adjacent nights falling on different weekdays. While autocorrelation is generally weak, there is a time-dependent structure within the data that should be considered when the data are modeled.

When data are analyzed by day of the week (Figure 19), most subjects show relative stability in their Sleep Duration, albeit with some inter-individual variation: S6 has the longest sleep with a low variability on Wednesdays, while S4 has the longest sleep on Saturdays with gradually decreasing sleep until Tuesdays. Similarly, some individuals (S2 and S6) report stable sleep quality ratings during the week, while the others rate their weekend sleep better.

**Figure 19.**
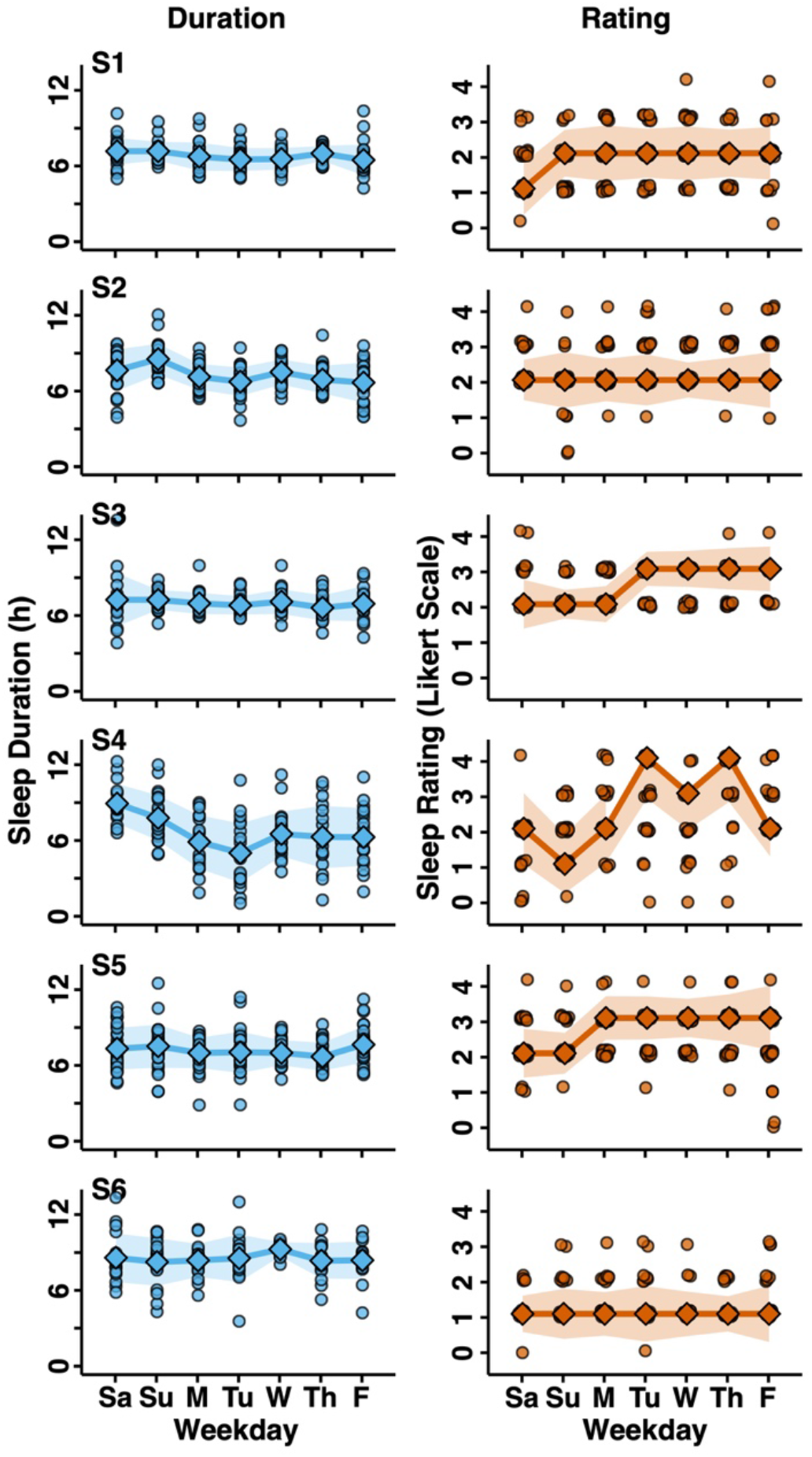
Weekday patterns of Sleep Duration and Sleep Rating. Sleep Duration and Sleep Rating are plotted separately for each weekday in each of the six subjects of Study 1 that vary depending on the day of the week for some subjects. While some subjects show relative stability in Sleep Duration (S1, S3, S5), others show strong effects of weekday (S2, S4). The self-report Sleep Rating is also variable with a tendency for the nights before the weekend days to be rated better (Saturday, Sa; Sunday, Su). Circles representing data points for Sleep Rating are jittered for visualization. Means (for Sleep Duration) and medians (for Sleep Rating) are shown by enlarged symbols with the shaded surround representing the standard error of the estimate.

We use a circle plot to show the sleep onset and offset distributions for each subject separately by weekday (Figure 20); this type of plot allows the sleep patterns to be visualized comprehensively on an intuitive mapping. Each point indicates the beginning or end of the major Sleep Episode and their medians. While there are subjects with stable average sleep onset (S1) and offset (S6), the plots confirm different effects of weekdays for different subjects, such as the effect of weekend from Friday to Monday on late sleep of S5, the effect of Monday, Tuesday, and Thursday on very late sleep of S4, and the effect of Monday, Tuesday, and Thursday on earlier wake up of S2. That is, specific structured sleep patterns are highly idiosyncratic to the individual.

**Figure 20.**
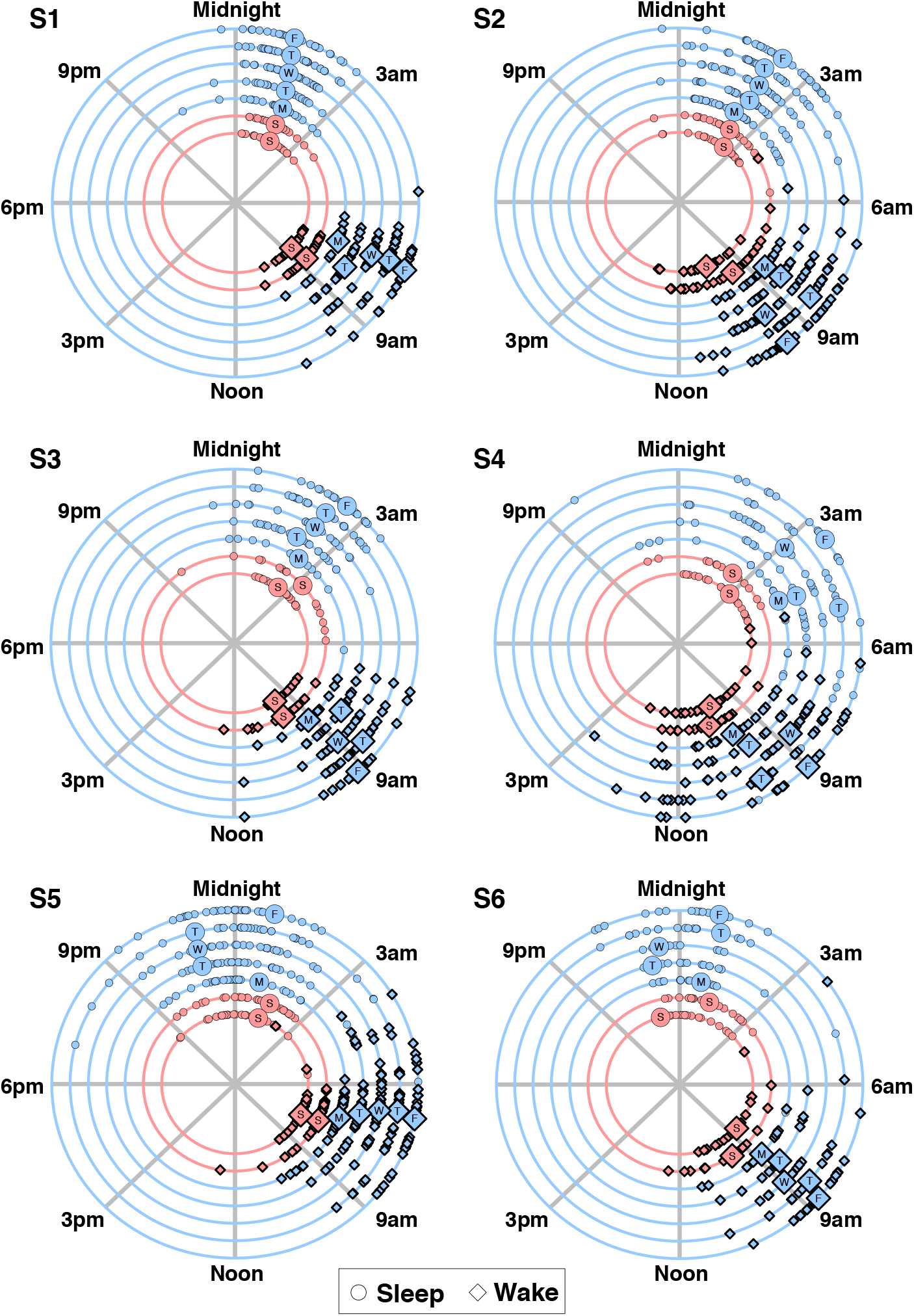
Weekday patterns of sleep onset and sleep offset. Sleep onset and offset (wake) times show differences depending on the day of the week. Each subject’s sleep onset (circles) and offset (diamonds) times are displayed (large circles/diamonds = medians) on a 24-hour clock format. Between-subject sleep onset and offset times are variable, as well are weekday patterns within the same individuals. These structured patterns, which can vary from subject to subject, should be considered when the data are modeled.

### Sleep Duration and Sleep Ratings Track Caffeine Intake in Some Individuals

Daily caffeine consumption is also assessed. Figure 21 shows the variation in caffeine consumption ranging from 0 (none) to 4 (5+ drinks) for each of the six subjects in Study 1; Figure 22 shows the weekly variation; Figure 23 shows the individual-level linear model for each subject modeling the relationship between caffeine consumption and Sleep Duration. While the effect is quantitatively small, four of the six subjects show a statistically significant relationship such that more caffeine consumption predicts a shorter Sleep Duration for the upcoming night.

**Figure 21.**
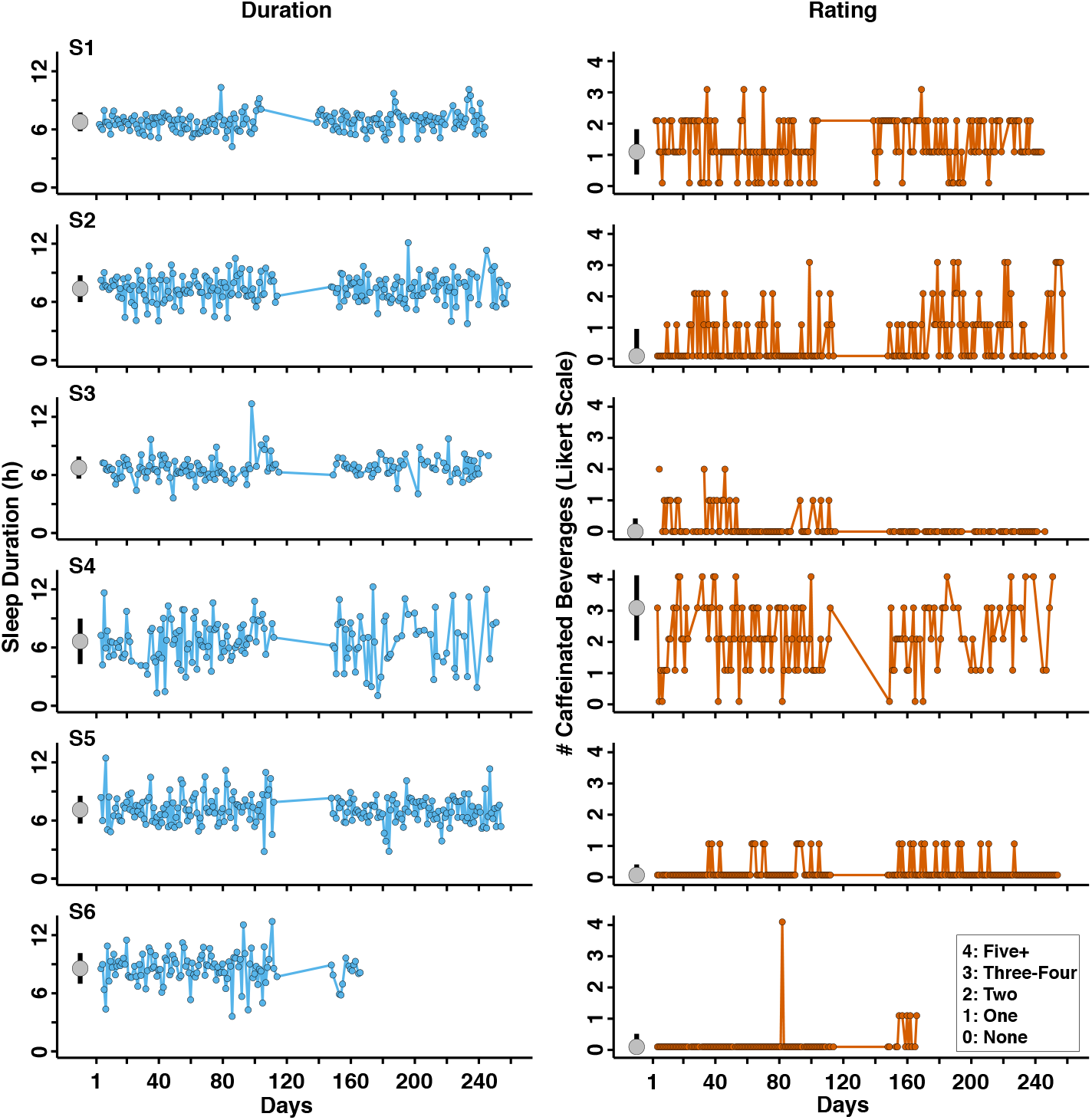
Longitudinal Behavior of Sleep Duration and caffeine consumption. The time course of each subject’s actigraphy-based Sleep Duration (in minutes) is plotted (left column) next to the same subject’s self-report caffeine consumption (right column; 5-point Likert scale; 0=None, 4=Five+). The plot format is similar to Figure 16. Even in this small sample, the between-subject and within-subject variability in caffeine consumption is notable, with certain subjects drinking caffeinated beverages often and variably across days (S1) and others consuming minimal caffeine on average with exceptions (S6).

**Figure 22.**
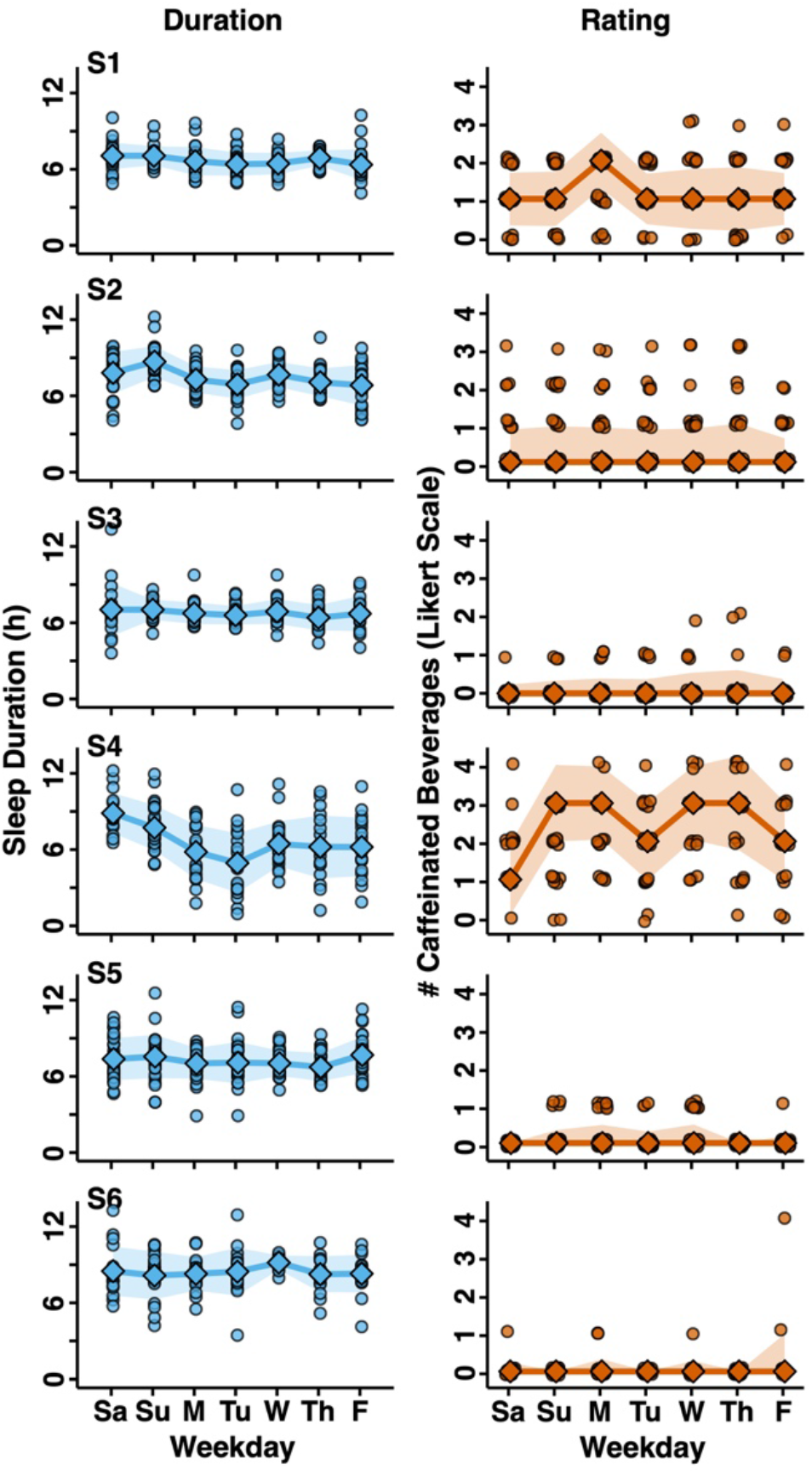
Weekly structure of Sleep Duration and caffeine consumption. Sleep Duration and caffeine consumption plotted separately for each weekday in each of the six subjects of Study 1 show variation depending on the day of the week. Circles representing data points for # Caffeinated Beverages are jittered for visualization. Means (Sleep Rating) and medians (# Caffeinated Beverages) are shown by enlarged symbols with the shaded surround representing the standard error of the estimate.

**Figure 23.**
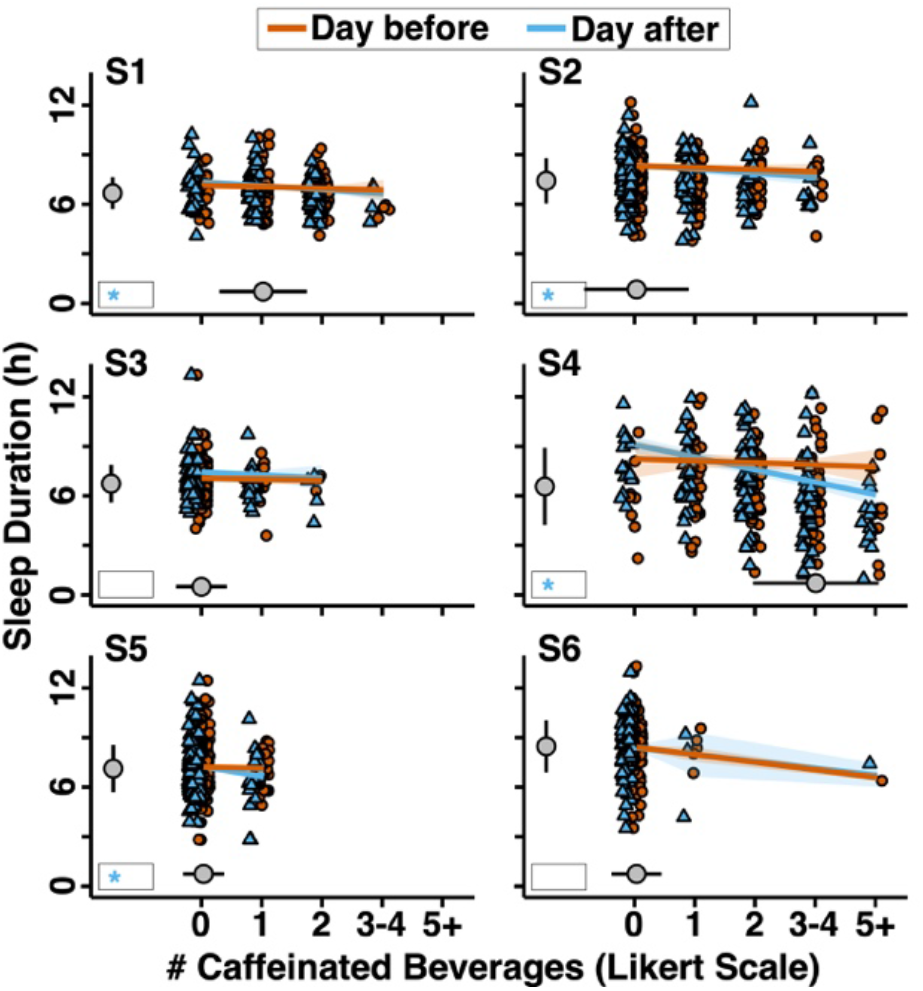
Self-report caffeine consumption predicts Sleep Duration. The individual-level linear model framework shows individualized associations between Sleep Duration and self-report ratings of the number of Caffeinated Beverages consumed. Data are plotted similar to Figure 17 and illustrate significant associations in certain subjects (S1, S2, S4, S5) but not others (S3, S6).

### Example Use Case: Sleep Patterns Show State Variation in Severe Mental Illness

To explore the feasibility and utility of sleep measures from extensive longitudinal assessments in patients, data from two individuals managing severe mental illness from Study 2 are analyzed using the DPSleep pipeline. Individuals participated for 543 and 309 days, with 89% and 59% of completed days of data obtained after data loss due to missingness and quality control.

Figure 24 illustrates data from S11 who is managing major depression. Of interest is the slowly changing sleep patterns that can be immediately visualized in the activity score plots (Figure 24A). Two separate features of the data are interesting and require distinct measures to quantify. The first is that the shifts to low-level activity, indicative of sleep, begin earlier and end considerably later across two long episodes that begin near Day 110 and Day 300. The reduced activity scores extend until noon on many days. The change in sleep onset and sleep offset and increase in the Sleep Duration is quantified with a 14-previous-days sliding window with less than three missing values tolerance in the DPSleep output shown in Figure 24C. The Sleep Timing Regularity Index (STRI) is also calculated as a 0-1 similarity index for 24-hours sleep and wake minutes of every day compared to an assumed day with the averaged sleep onset and offset. The STRI represents a modified version of the Sleep Regularity Index (SRI) introduced by Phillips and colleagues (2017). The SRI “calculates the percentage probability of an individual being in the same state (asleep vs. awake) at any two minutes 24 hours apart”, thus focusing on day-to-day, circadian fluctuations in sleep. The STRI, our modified version of the SRI, compares each study day to each participant’s “average sleep day”.

**Figure 24.**
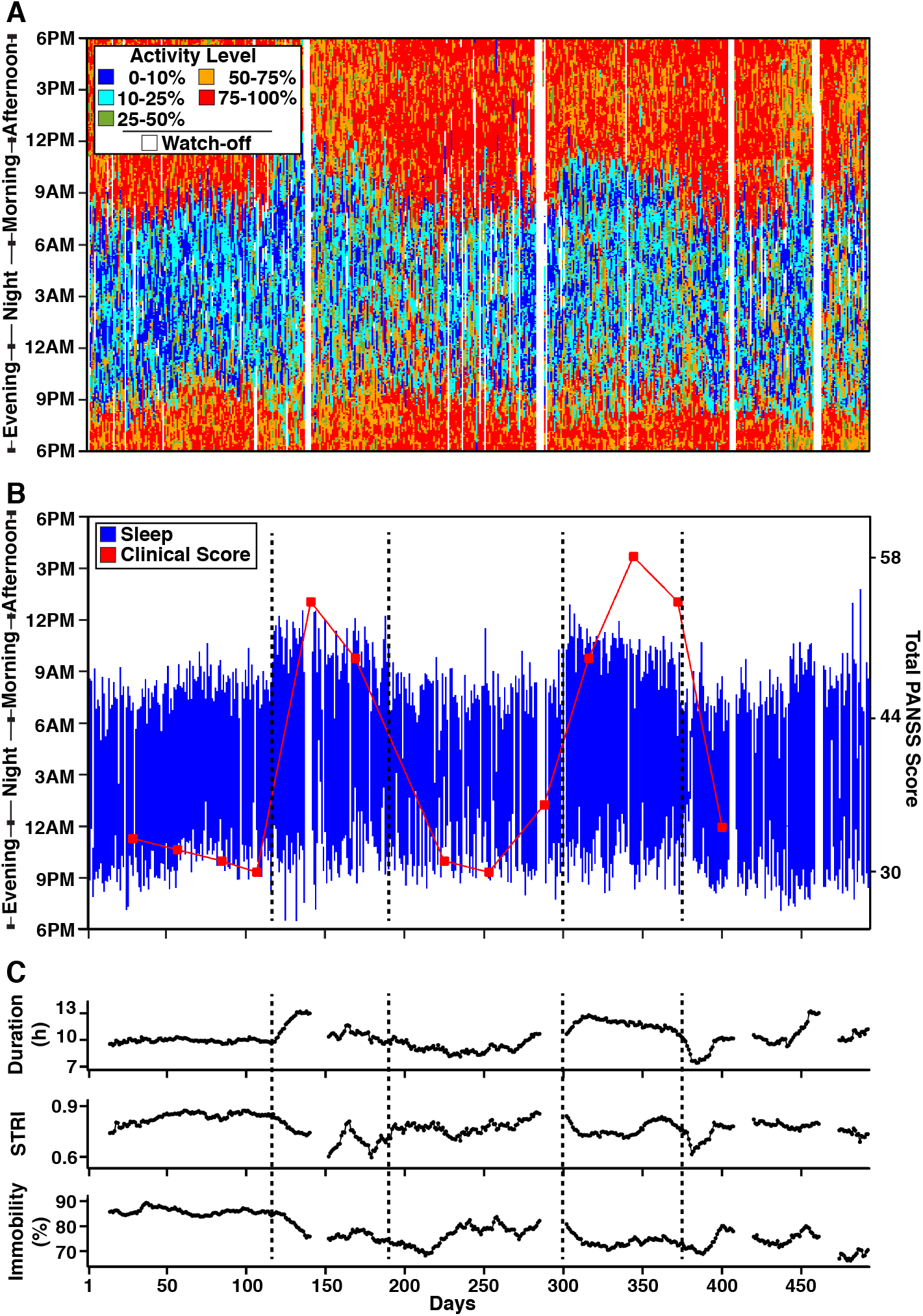
Example longitudinal sleep pattern over 500 days in a patient with severe mental illness. Three panels display longitudinal Activity Score, Sleep Episode estimates and clinical severity score, and quantitative metrics derived from the actigraphy for S11 of Study 2. (A) The top panel displays the continuous Activity Scores colored by the threshold for each day, similar to Figure 7A. (B) The middle panel displays the time-zone adjusted and fully quality-controlled estimates of the Sleep Episodes (similar to Figure 7C) with the severity of the Clinical Score overlaid by a red line. The Clinical Score reflects the total score on the Positive and Negative Syndrome Scale (PANSS). (C) Temporally-smoothed (14-day backward moving average) estimates of three activity-based measures are plotted: the estimated Sleep Duration (labeled Duration), the Sleep Timing Regularity Index (labeled STRI), and the *SleepImmobility* percentage (labeled Immobility). Gaps in the plots reflect missing days; if more than two days were missing, the temporal average that would include those days are absent. Clear state changes in the sleep patterns can be observed (demarcated by dashed black lines in B) that are temporally coincident with negative changes in the Clinical Score.

Computing a participant’s average sleep day involves several steps to select the minutes from each day during which a participant is most frequently asleep such that the total duration is equal to the participant’s average daily Sleep Duration. The daily STRI is then computed by comparing each daily Sleep Episode to the average Sleep Episode, at the minute-level, and computing the proportion of minutes for which these two Sleep Episodes matched in sleep. This value, demonstrated in Figure 24C with a backward 14-day sliding window, shows noticeable drops that roughly occur at the beginning of the period of longer sleep intervals. Data with more than 3 missing values during the sliding window are considered missing.

What is further notable is that the major Sleep Episodes include higher activity score periods than during typical sleep, suggesting an interrupted and inconsistent sleep. This feature is picked up in the derived measures of *SleepImmobility* percentage, defined in Table 1 as the percentage of *ImmobileMinutes* in the major Sleep Episode illustrated as similar sliding window values in Figure 24C. Adopting a previously described quantitative framing of Immobility (version 7.2, CamNtech, Cambridge, UK, 2008; Meltzer et al., 2015), the *ImmobileMinutes* were defined in Table 1 as minutes with lower than a cut-off threshold (40th percentile activity here) that is the threshold to show the Activity Level in Figure 4. To further illustrate the utility of these data, the clinical severity of illness, measured via the Positive and Negative Syndrome Scale (PANSS) Total Score, is shown overlaid on top of the major Sleep Episode in Figure 24B. The periods of sleep disruption and extended sleep offsets correspond to the periods of high illness severity.

Figure 25 illustrates data from S12 who lives with psychosis associated with schizophrenia. The activity scores display an irregular and slowly drifting pattern, including a shift to low activity episodes that occur later in the day around Day 55 and then an earlier shift beginning near Day 70 (Figure 25A). The period of most severe illness symptoms occurs near to the most irregular sleep periods after Day 75. Note also this individual has generally poorer clinical scores (Figure 25B) as compared to S11 and a generally lower STRI (Figure 25C). These data illustrate the complexity and richness of information that can be obtained through extensive longitudinal analysis of the actigraphy data, and also how different each individual can be from one another.

**Figure 25.**
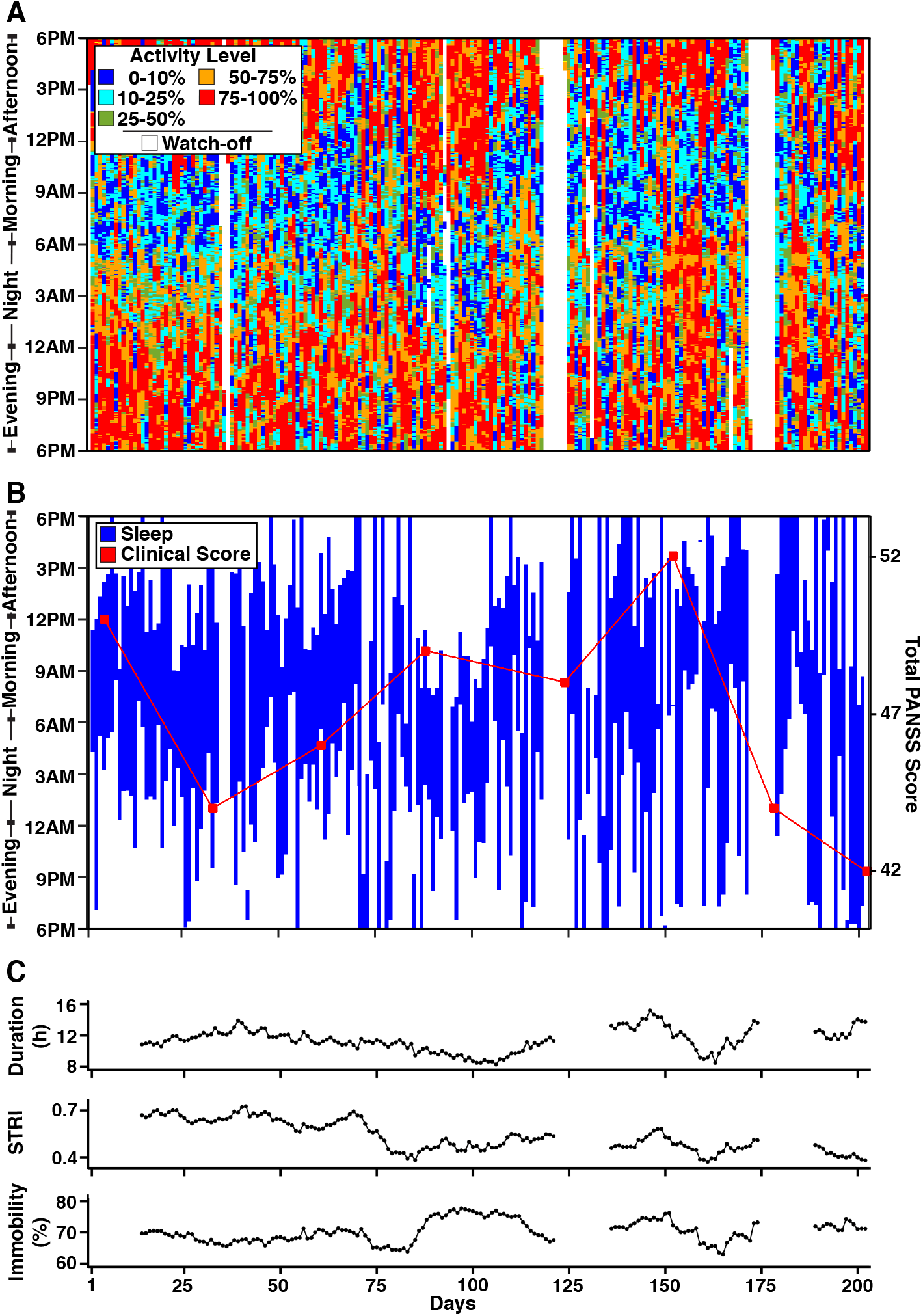
Example longitudinal sleep pattern over 200 days in a patient with severe mental illness. Using the same format as Figure 24, data from S12 of Study 2 are plotted. Note the consistently irregular and fragmented sleep patterns that are pervasive throughout the course of the measurements. These data illustrate the feasibility of collecting high-quality continuous data using passive actigraphy in individuals with severe mental illness.

### Example Use Case: Sleep Patterns in Relation to Deep Dynamic Phenotyping

To demonstrate how sleep measures can be combined with additional forms of digital phenotyping information, two individuals from Study 1 are displayed in Figures 26 and 27. In each plot, the sleep patterns for each individual are illustrated through the course of an academic year, along with the quantified measures of Sleep Duration and STRI. In addition, self-report measures of social and academic behaviors (time on Homework and time Interacting) and mood (Happy and Stress) are displayed aligned to the estimates of the longitudinal Sleep Episodes. The examples illustrate clear dynamics of sleep, including weekend (empty circles) / weekday (filled circles) effects, changes between the active semester and breaks, as well as intermittent deviations for regular patterns. These data illustrate the potential of using low-burden wearables in combination with smartphone-based digital phenotyping to capture a great deal of information about life rhythms and changes due to environmental demands.

**Figure 26.**
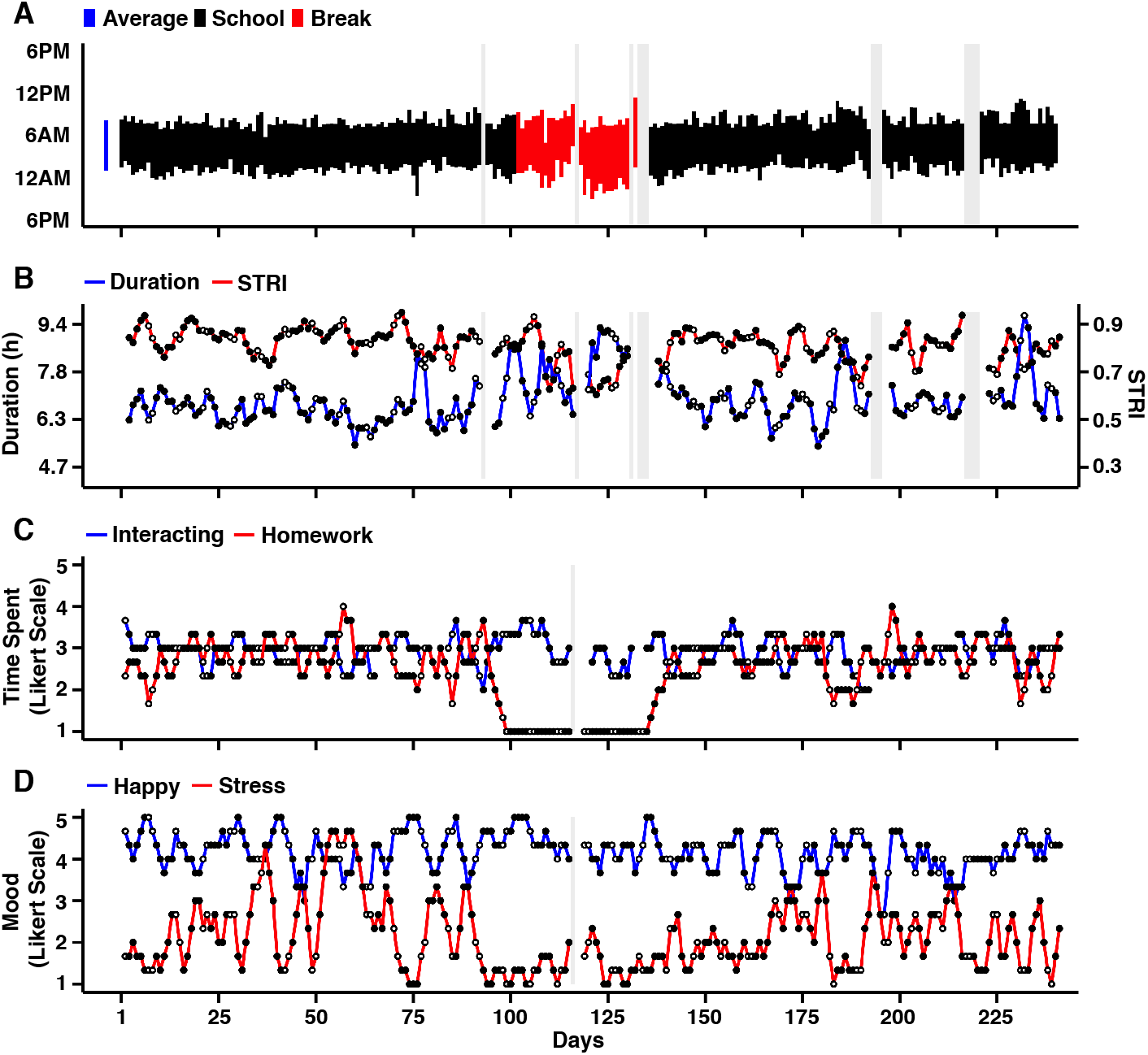
Example year in the life of college student S1. Four panels display multi-modality longitudinal data from the full academic year in a college undergraduate (S1 of Study 1). (A) Actigraphy-based estimates of the Sleep Episode are displayed, colored by academic period (black = academic school year; red = winter break). Missing data with gray backgrounds reflect missing (wrist-off) or quality removed data (e.g., on a travel day across time zones). The mean Sleep Episode is shown to the left in blue. (B) Temporally-smoothed (3-day backward moving average with no missing tolerance) estimated of the Sleep Duration (Duration) and Sleep Timing Regularity Index (STRI). (C) Temporally-smoothed estimates of time Interacting and time doing Homework from the self-report questionnaire (see Appendix I). (D) Self-report estimates of mood, including happiness (Happy) and stress (Stress) (also see Appendix I). This individual displays stable and regular sleep patterns with periodic deviations. Note the deviations in time Interacting and Homework during the break period.

**Figure 27.**
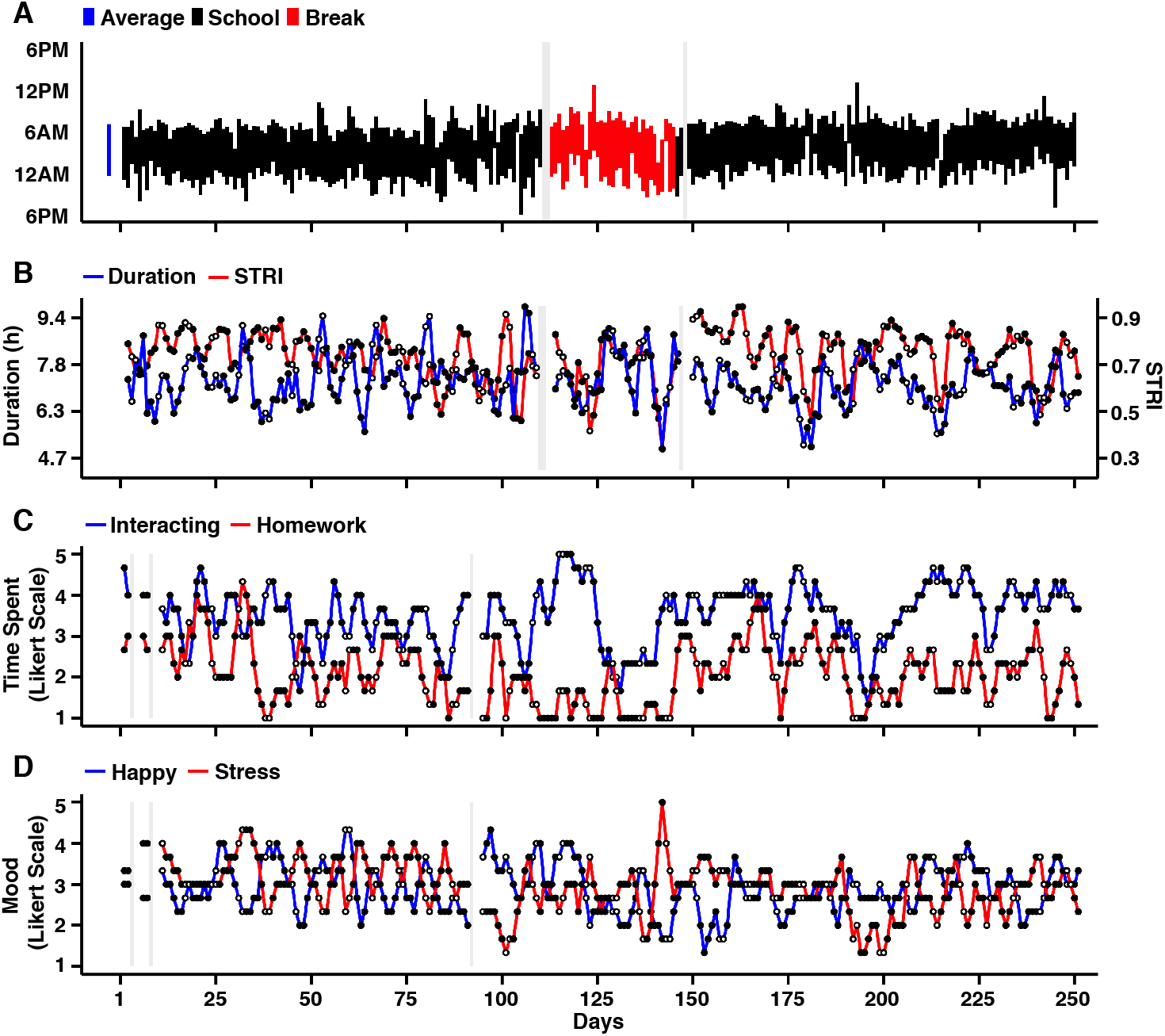
Example year in the life of college student S5. Using the same format as Figure 26, four panels display multi-modality longitudinal data from the full academic year in a college undergraduate (S5 of Study 1). This individual shows highly irregular sleep during the academic year and during the break. Note that in addition to the general variation in sleep patterns and mood, there are specific instances of extreme variations that are reflected in Sleep (e.g., Day 144 in B) and Stress (D).

### An Example of Limitations

In showing examples of the utility of the DPSleep processing pipeline and example applications, we wanted to also show, as a last point, a clear example of a limitation. The present pipeline and quality control adjustments make assumptions that have been selected because they work most of the time in the range of contexts for which they were tested. But real-life happenings are complex. Figure 28 plots an interesting example. In this example, the daily data from Figure 3B is replotted along with phone use and GPS location data from the same participant. What is notable is that our original estimate, based only on watch actigraphy data and the button press, combined with the rule employed to join together episodes of low activity as a continuous major Sleep Episode, likely mischaracterizes this night’s true continuous Sleep Episode. What the phone data reveal is that the nighttime period of activity is followed by missing GPS data, intermittent phone use, and then eventual arrival at a new location. This individual most likely woke up early and got on a bus or a train. Our automated estimation procedures as well as our quality control, using only the watch actigraphy, mischaracterizes this Sleep Episode.

**Figure 28.**
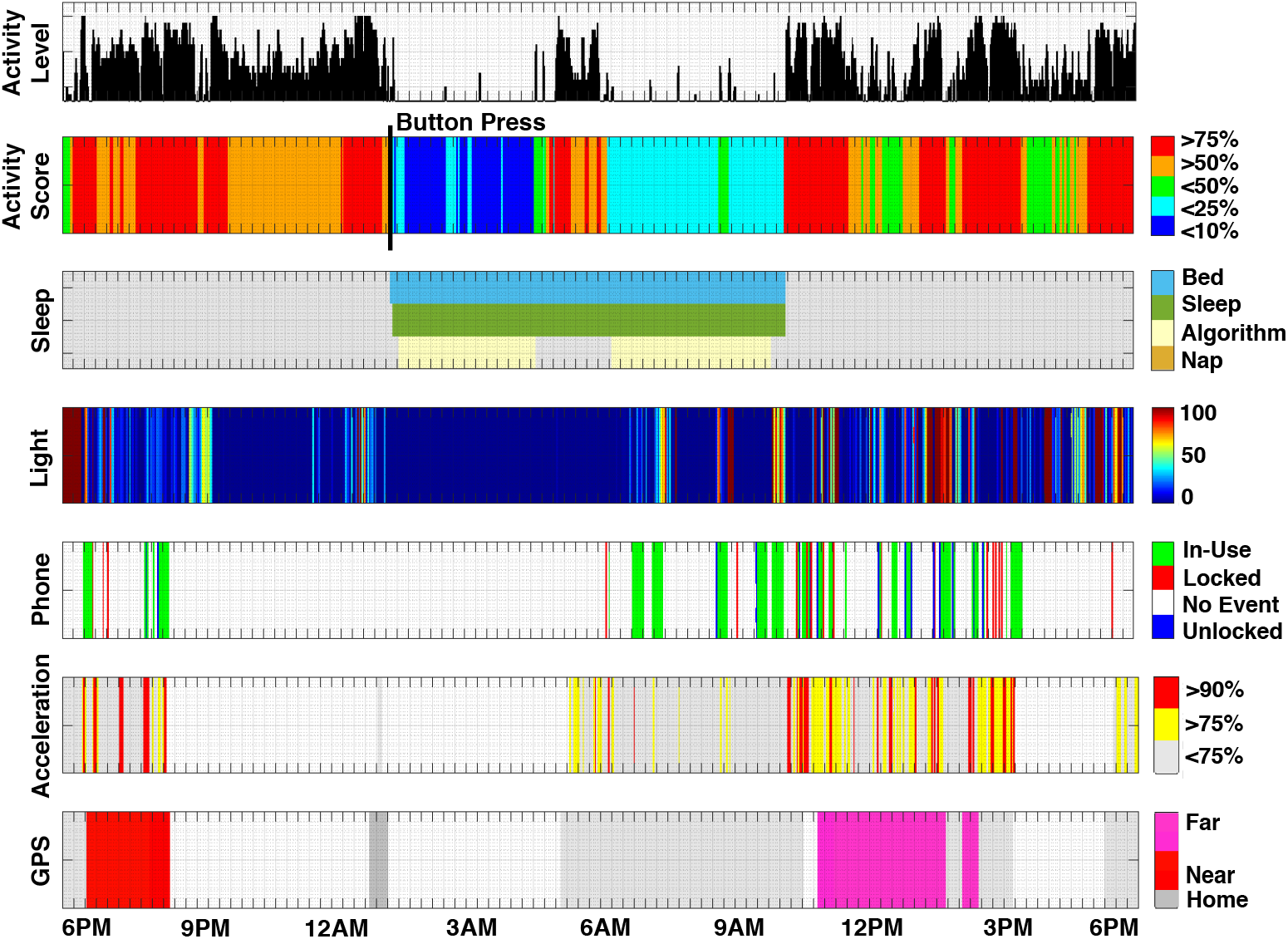
Example of an erroneous estimate of the Sleep Episode. Plots of a night from Study 1 illustrate that actigraphy-based sleep estimation is imperfect. The top four rows represent the watch-based actigraphy information as well as our present procedure’s estimates of the Sleep Episode (from Figure 3B). The bottom three rows show the phone use data for this night plotted in an aligned format. Note that the ‘active’ period near 5AM is followed by periods of phone use and acceleration, and then a GPS-estimated change in location (see pink location estimate starting at 11AM). This subject likely got out of bed and went into a car or bus, sitting relatively motionless for several hours, until she or he arrived at a new location. Our procedures, which only use the watch data, mischaracterized this Sleep Episode. This complex night is a reminder that real-world behavior is complex and that the assumptions of the approach are imperfect for all situations.

## Discussion

The present work describes and demonstrates the utility of an open-source, longitudinal sleep-analysis platform called DPSleep. The platform was applied to two cohorts of participants that possessed extended data over months to years and included clinically healthy undergraduates as well as outpatients living with severe mental illness. The goal of these diverse explorations was to validate the approach and demonstrate its utility across multiple real-world participant cohorts. The results revealed that the approach could capture the major Sleep Episode and detect dynamic patterns of sleep behavior including in individuals presenting with episodic clinical illness (e.g., Figures 24-25).

Several prior studies have investigated the validity and accuracy of wrist-worn actigraphy devices to estimate sleep parameters and compare them against gold-standard PSG-derived measures and self-report sleep quality (Khademi et al., 2019). DPSleep uses frequency-based analysis and begins with the high-frequency raw data from the wrist-worn accelerometer. The high precision of the sampling frequency and the continuity of the collected data in our samples provided ideal datasets to apply frequency-based analysis, a robust and efficient tool to analyze the activity of the individual, that complements alternative procedures such as zero crossing mode (ZCM), time above threshold (TAT), or digital integration mode (DIM) (Stone and Ancoli-Israel, 2017).

Frequency-based analysis inherently and automatically discounts the alternating gravity effect on different axes, without the need to eliminate the amplitude of the signal (van Hees et al., 2015), and at the same time disentangles the natural high frequency shakes of the body from the actual repositioning dynamic, which is the main focus in sleep research. As explained earlier, the raw accelerometer data contains the gravity component offset, that can be removed automatically in frequency analysis when focusing on higher (>0) frequencies. A challenge with our approach was related to participant burden as the participants had to visit the lab every 4-5 weeks to get a refreshed memory and battery to prevent data missingness. An alternative solution to attenuate the burden on subjects is to use wireless data transfer and cloud storage; however, this approach has its own limitations, based on the wireless storage size and battery usage in addition to the data security concerns. We expect advances in widely available commercial technologies to gradually alleviate these current existing challenges.

In our study, reported sleep quality shows a strong correlation with the activity-based Sleep Duration measure; therefore, the focus of our sleep algorithm development was mainly to optimize this parameter, meanwhile building a platform to explore other relevant sleep-related parameters in future work. Despite existing sleep detection algorithms that look for 5-15 low activity minutes (van Hees et al., 2015; Hees et al., 2018), DPSleep starts with structural analysis of the daily activity to first detect the large episode of the day with lowest average activity; and then the edges of the episode are adjusted using smaller moving windows and heuristic rules. This approach is an efficient solution to eliminate the need for any kind of sleep diary or ambient light assumptions due to the fact that the former is not usually available and accurate, and the later could be misleading, since the wristband can be blocked at any time of the day by a long sleeve. Moreover, this approach distinguishes the short naps or inactive periods during the day from the long, continuous, though disrupted, Sleep Episodes without any assumptions about the sleep time. Structural analysis of the activity scores during the whole study, using the individual’s statistics and moving average windows with different sizes, suggests the most likely episode for the individual’s sleep during the day. Beside the moving average windows to automatically detect the major Sleep Episode, DPSleep provides a helpful tool to the investigators to decide, with reasonable confidence, about idiosyncratic sleep behaviors such as no sleep or very short Sleep Episodes. Additional button presses, if available, and adjacent activities are used to tune this episode and/or connect smaller pieces to shape the entire Sleep Episode.

### Caveats and Limitations

As we illustrate in Figure 28, our approach can make mistakes. In this sense, we illustrate how inexpensive and easy-to-obtain actigraphy data can be analyzed to estimate the major Sleep Episode and dynamic patterns over many days. However, the real world is messy. Atypical behavioral patterns (e.g., many short naps without a clear extended primary Sleep Episode) and behaviors that yield extreme accelerometer readings are challenging for our approach. Specifically, using our methods, actigraphy-based sleep detection is not appropriate for measuring sleep when the individual is on a shaking platform such as a plane, train, or bus. This is an unavoidable situation in longitudinal studies; however, since the on-plane sleep effect was not the focus of our study, and those days were very few compared to the days of the entire study, we were able to detect those days using GPS data and eliminate them from the results. To solve the continuous shaking challenge in similar situations or in studies on individuals with Parkinson’s disorder, sleep detection devices based on ambulatory circadian monitoring (ACM) are recommended (Madrid-Navarro et al., 2019). Most broadly, as experience with actigraphy grows and repositories of annotated data become available from common and less common activities, the present approach (and others) can be refined to handle a wider variety of situations. The validation and exploration of DPSleep, as illustrated in these initial explorations, provides a tool that can be used today as well as refined and expanded through its open-source release.

## Acknowledgements

We thank Laura Farfel, Marisa Marotta, Erin Phlegar, Lauren DiNicola, Arpi Youssoufian, Elyssa Barrick, Nora Mueller, Crystal Blankenbaker and Joanna Tao, for their help collecting data, and Elizabeth Klerman for valuable comments and edits on the manuscript. Timothy O’Keefe, Harris Hoke, Lily Jeong and Sarah Guthrie provided valuable assistance in neuroinformatics support. Kenzie W. Carlson helped with the Beiwe component of the studies. Katherine Miclau, Amira Song, Emily Iannazzi and Hannah Becker helped with the quality control procedure. Patrick Mair provided valuable consultation on statistical approaches. Work was funded by a generous gift from Kent and Liz Dauten, NIMH grants P50MH106435 and U01MH116925, and NIH award DP2MH103909.

## Appendix I Self-Report Questionnaire

### Sleep

How did you sleep last night?

1. Terribly: little or no sleep
2. Not so well: got some sleep but not enough
3. Sufficient: got enough sleep to function
4. Good: got a solid night’s sleep and felt well-rested
5. Exceptional: one of my best nights of sleep

### Caffeine

In the past 24 hours how much caffeine (e.g., 8oz of coffee, tea, or soft drink) did you consume?

1. None
2. One drink
3. Two drinks
4. Three or four drinks
5. Five or more drinks

### Interacting

How much of your waking time did you spend interacting with others over the past 24 hours?

1. Very little of my time (0-20%)
2. Some of my time (21-40%)
3. About half of my time (41-60%)
4. Most of my time (61-80%)
5. Almost all of my time (81-100%)

### Homework

How much of your waking time did you spend studying or doing homework over the past 24 hours?

1. Very little of my time (0-20%)
2. Some of my time (21-40%)
3. About half of my time (41-60%)
4. Most of my time (61-80%)
5. Almost all of my time (81-100%)

### Happy

How much did you feel happy over the past 24 hours?

1. Very slightly or not at all
2. A little
3. Moderately
4. Quite a bit
5. Extremely

### Stress

How much did you feel stressed over the past 24 hours?

1. Very slightly or not at all
2. A little
3. Moderately
4. Quite a bit
5. Extremely

## Appendix II Quality Control Instruction

On the <u>Activity Score plot</u>: The activity level in each minute is color-coded so that Cyan and Blue are the <u>lowest activity</u> minutes and Orange and Red are the minutes with the <u>highest activity</u>. Green is the <u>medium activity</u>. The Sleep Episode is almost continuous appearance of Blue and/or Cyan (lowest activity).

On the <u>Sleep plot</u>:

- **Light yellow box** is the initial provisional estimate of the Sleep Episode automatically detected by the algorithm (called Algorithm on the Sleep plot).
- **Green box** is the <u>Sleep Episode</u> estimation when the sleep edges are adjusted to:
  1. Button Press Markers: if the marker(s) exist(s) and is(are) less than <u>60 minutes</u> inside the sleep box detected by the algorithm; otherwise:
  2. <u>When the edge of the Algorithm box is within the Cyan/ Blue block:</u> Choose the first minute out of the Algorithm box that does not have the lowest activity. It can be Green, Orange or Red. In the other word, extend the Sleep Episode to the edge of the continuous Blue/Cyan block.
  3. <u>When the edge of the Algorithm box is out of the Cyan/ Blue block:</u> Choose the first minute inside the Algorithm box that has the lowest activity. In other words, shrink the Sleep Episode to the edge of the continuous Blue/Cyan block.
- **Blue box** is the <u>Bedrest Episode</u> estimation which is set to:
  1. Button Press Markers: if the marker(s) exist(s) and is(are) less than <u>60 minutes</u> outside the sleep box detected by the algorithm; otherwise:
  2. When the edge of the Algorithm box is within a Cyan/ Blue block: Choose the first minute out of the Algorithm box that has high activity. It can be Orange or Red, but not Green. In other words, extend the Bedrest Episode to the first high active minute.
  3. When the edge of the Algorithm box is out of the Cyan/ Blue block: Choose the first minute outside the <u>Sleep Episode</u> box that has high activity (Orange or Red).

**Notice:** In case there are short intervals of detected watch-off minutes (White) during sleep, which are not larger than 150 minutes consider them as very low activity minutes (Blue). These are the very low activity minutes occasionally happening during the Sleep Episode that the sliding window erroneously tags as watch-off, and the activity of the minutes around them shows that the watch is actually not off the wrist and they are part of the Sleep Episode.

**Notice:** If there is no Sleep Episode that contains more than 45 <u>continuous lowest activity</u> minutes, report <u>No Sleep</u>.

**Notice:** The general rule for the <u>Sleep Episode</u> is to choose the longest episode that contains one or more low activity epochs, and multiple epochs can be connected to each other with no more than 90-min active epochs (bouts) in between, and choose the rest as <u>Nap Periods</u>. If the longest epoch is after 12PM and there is a shorter epoch before, choose the shorter one as <u>Sleep Episode.</u>

